# Data-driven beamforming techniques to attenuate ballistocardiogram (BCG) artefacts in EEG-fMRI without detecting cardiac pulses in electrocardiography (ECG) recordings

**DOI:** 10.1101/2020.11.27.401851

**Authors:** Makoto Uji, Nathan Cross, Florence B. Pomares, Aurore A. Perrault, Aude Jegou, Alex Nguyen, Umit Aydin, Jean-Marc Lina, Thien Thanh Dang-Vu, Christophe Grova

**Author notes:** **Corresponding authors**: Concordia University, 7141 Sherbrooke St. West, SP 165.30, Montreal H4B 1R6, Canada., E-mail addresses (M. Uji), (C. Grova).

## Abstract

Simultaneous recording of EEG and fMRI is a very promising non-invasive neuroimaging technique, providing a wide range of complementary information to characterize underlying mechanisms associated with brain functions. However, EEG data obtained from the simultaneous EEG-fMRI recordings are strongly influenced by MRI related artefacts, namely gradient artefacts (GA) and ballistocardiogram (BCG) artefacts. The GA is induced by temporally varying magnetic field gradients used for MR imaging, whereas the BCG artefacts are produced by cardiac pulse driven head motion in the strong magnetic field of the MRI scanner, so that this BCG artefact will be present when the subject is lying in the scanner, even when no fMRI data are acquired. When compared to corrections of the GA, the BCG artefact corrections are more challenging to remove due to its inherent variabilities and dynamic changes over time. Typically, the BCG artefacts obscure the EEG signals below 20Hz, and this remains problematic especially when the frequency of interest of EEG signals is below 20Hz, such as Alpha (8-13Hz) and Beta (13-30Hz) band EEG activity, or sleep spindle (11-16Hz) and slow-wave oscillations (<1 Hz) during sleep. The standard BCG artefact corrections, as for instance Average Artefact Subtraction method (AAS), require detecting cardiac pulse (R-peak) events from simultaneous electrocardiography (ECG) recordings. However, ECG signals in the MRI scanner are sometimes distorted and will become problematic for detecting reliable R-peaks. In this study, we focused on a beamforming technique, which is a spatial filtering technique to reject sources of signal variance that do not appear dipolar in the source space. This technique attenuates all unwanted source activities outside of a presumed region of interest without having to specify the location or the configuration of these underlying source signals. Specifically, in this study, we revisited the advantages of the beamforming technique to attenuate the BCG artefact in EEG-fMRI, and also to recover meaningful task-based induced neural signals during an attentional network task (ANT) which required participants to identify visual cues and respond as accurately and quickly as possible. We analysed EEG-fMRI data in 20 healthy participants when they were performing the ANT, and compared four different BCG correction approaches (non-BCG corrected, AAS BCG corrected, beamforming+AAS BCG corrected, beamforming BCG corrected). We demonstrated that beamforming BCG corrected data did not only significantly reduce the BCG artefacts, but also significantly recovered the expected task-based induced brain activity when compared to the standard AAS BCG corrections. Without detecting R-peak events from the ECG, this data-driven beamforming technique appears promising especially for longer data acquisition of sleep and resting EEG-fMRI. Our findings extend previous work regarding the recovery of meaningful EEG signals by an optimized suppression of MRI related artefacts.

**Highlights:**

- Beamforming spatial filtering technique attenuates ballistocardiogram (BCG) artefacts in EEG-fMRI without detecting cardiac pulses in electrocardiography (ECG) recordings.
- Beamforming BCG denoising technique recovers expected task-based induced visual alpha and motor beta event-related desynchronization (ERD).
- Beamforming technique improves signal-noise ratios (SNR) of neural activities as compared to sensor level signals.
- Data-driven beamforming technique appears promising for longer data acquisition of sleep and resting EEG-fMRI without relying on ECG signals.

## Introduction

Electroencephalography (EEG) and functional magnetic resonance imaging (fMRI) are two non-invasive neuroimaging techniques that are often used to investigate human brain function, so that their multimodal integration has been actively considered (Jorge et al., 2014; Laufs, 2012; Moeller et al., 2020; Murta et al., 2015). fMRI measures hemodynamic Blood Oxygen Level Dependent (BOLD) responses driven by increased metabolic demands during neuronal activation, which means indirect measurements of brain activity (Logothetis et al., 2001), whereas EEG measures the electrical potentials generated by the cooperative action of neurons providing a more direct measure of neuronal activity (Buzsáki et al., 2012). Furthermore, fMRI has a high spatial resolution (in order of mm) with a poor temporal resolution (in order of seconds) due to slow hemodynamic responses, while EEG generally has a high temporal resolution (in order of milliseconds) with a poor spatial resolution. Simultaneous recordings of EEG and fMRI are very promising non-invasive techniques, provide a wide range of complementary information, and can be advantageous for improving our understanding of human brain function.

Simultaneous EEG-fMRI has been successfully used to investigate epileptic networks in patients with drug-resistant focal epilepsy undergoing pre-surgical evaluation (Gotman et al., 2006; Gotman and Pittau, 2011; Grova et al., 2008; Heers et al., 2014; Ives et al., 1993; Lemieux et al., 2001; LeVan and Gotman, 2009; Murta et al., 2012), and sleep (Cross et al., 2021; Dang-Vu et al., 2011, 2008; Fultz et al., 2019; Hale et al., 2016) enabling spatial localizations of the hemodynamic responses elicited by temporally dynamic specific EEG features/events. Recently, the application of EEG-fMRI has been extended to investigate the spatiotemporal dynamics of neural activity (for a review, see Huster et al., 2012), and also to study the underlying neurophysiological origins of the measured responses by comparing neural and hemodynamic signals (Mullinger et al., 2013). The primary advantage of simultaneous EEG-fMRI acquisition over separate recordings is that it enables the investigation of unpredictable or spontaneous brain activity, and the study of the trial-by-trial covariation in brain processing as measured by the two techniques (Bagshaw et al., 2004; Becker et al., 2011; Debener et al., 2006; Eichele et al., 2008; Goldman et al., 2002; Horovitz et al., 2008; Mayhew et al., 2013; Mobascher et al., 2009; Mullinger et al., 2014; Olbrich et al., 2009; Scheibe et al., 2010; Uji et al., 2018). EEG-fMRI analysis has demonstrated specific BOLD correlates of distinct neurophysiological components including the auditory oddball (Bénar et al., 2007; Eichele et al., 2005) and the error-related negativity (Debener et al., 2005), as well as specific neural activity in specific frequency bands (Goldman et al., 2002; Laufs et al., 2003; Tyvaert et al., 2008). These studies have demonstrated that simultaneous EEG-fMRI can provide greater specificity and sensitivity when compared to a single neuroimaging modality (EEG or fMRI alone) regarding the spatial arrangement (Goldman et al., 2009; Novitskiy et al., 2011) and the temporal sequence (Eichele et al., 2005; Mayhew et al., 2012) of brain responses.

However, EEG data obtained from the simultaneous EEG-fMRI recordings are strongly influenced by MRI related artefacts (Allen et al., 2000, 1998; Niazy et al., 2005), which are gradient artefacts (GA) and ballistocardiogram (BCG) artefacts. The GA is induced by temporally varying magnetic field gradients used for MR imaging causing large voltage fluctuations in the EEG traces through Faraday’s law of induction, 10-100 times larger when compared to EEG signals induced by brain signals (Allen et al., 2000), therefore completely obscuring the measured EEG signal. However, since it is generated by the fMRI sequence itself, such artefact is highly reproducible and removal of the GA can be accurately achieved by subtracting averaged GA waveforms measured at each EEG electrode across fMRI volume acquisitions from each corresponding electrode. This method entitled the Average Artefact Subtraction (AAS) approach, is efficient because of the very precise and repetitive characteristics of GA features induced by fMRI sequence (Allen et al., 2000). However, AAS requires a high sampling rate in the EEG to measure GAs with appropriate temporal resolution (typically 5kHz), whereas MR compatible EEG amplifier should offer a large dynamic range to avoid any saturation of the artefact. Moreover, accurate synchronisation between the MRI scanner and EEG clocks significantly improves the quality of GA corrections (Mandelkow et al., 2006; Mullinger et al., 2008b). However, even after correction, residual GAs still obscure small-amplitude neuronal signals, mainly in the high frequency range, typically above 20 Hz, (Mullinger et al., 2011, 2008b). In this context, the use of an accelerated fMRI sequence (i.e. sparse multiband fMRI acquisitions) can allow obtaining some MRI gradient free quiet periods during the paradigm. In those segments that provide GA free EEG periods, one can then assess specific EEG frequency of interest above 20Hz, as suggested in Uji et al. (2018) when studying the coupling between single trial variability in gamma frequency (55-80Hz) EEG signals and fMRI BOLD responses during a motor task. In general, the conventional AAS approach remains the most reliable GA correction method when implementing with appropriate hardware solutions (synchronization: Mandelkow et al., 2006; Mullinger et al., 2008) and optimal head positioning (Mullinger et al., 2011).

Compared to the GA, BCG artefacts are definitely more challenging to cope with due to their inherent variabilities and dynamic changes over time (Laufs et al., 2008). The BCG artefacts are produced by cardiac pulse driven head motion in the strong magnetic field of the MRI scanner (Debener et al., 2008; Mullinger and Bowtell, 2011), so that this artefact will be present even when no fMRI acquisitions are performed. Typically, the BCG artefacts obscure the EEG signals below 20Hz (Bonmassar et al., 2002; Xia et al., 2014, 2013), and this remains problematic especially when the frequency of interest of EEG signals is below 20Hz, such as Alpha (8-13Hz) and Beta (13-30Hz) band EEG activity, or sleep spindle (11-16Hz) and slow-wave oscillations (<1 Hz) during sleep. In order to overcome this complexity, both software (Independent Component Analysis (ICA): Debener et al., 2007; Optimal Basis Set (OBS): Niazy et al., 2005) and hardware (Reference layer artefact subtraction: Chowdhury et al., 2014; Carbon-wire loop: Masterton et al., 2007; Optical Motion tracking system: LeVan et al., 2013) based solutions have been suggested for the BCG artefact corrections. In the software solutions, the ICA technique has been used to remove independent components (ICs) representing the BCG artefacts, whereas the OBS approach has used principal component analysis (PCA) to remove the principal components (PCs) representing the BCG artefacts. Although both ICA (Debener et al., 2008; Mantini et al., 2007; Srivastava et al., 2005) and PCA (Niazy et al., 2005) approach have successfully removed the BCG artefacts, objective and accurate classifications/selections of BCG-related ICs and PCs still remains a major concern (Abreu et al., 2016; Vanderperren et al., 2010). Furthermore, the ICA and PCA approaches have usually been applied after the AAS BCG approach to remove residual BCG artefacts (Abreu et al., 2018). Although these different methods have been suggested for the BCG artefact corrections using either software or hardware-based solutions, the most commonly used approach is still AAS approach (see a review Abreu et al., 2018). However, this AAS approach requires high precisions of cardiac pulse (R-peak) event detections in the MRI scanner which are used for subtracting averaged BCG artefact templates. R-peak events are detected from simultaneous electrocardiography (ECG) recordings. However, ECG signals in the MRI scanner are often distorted, which makes automatic detection of R-peaks problematic (Chia et al., 2000; Mullinger et al., 2008b), and is significantly time-consuming when manual correction is required. Detecting R-peak events from the ECG is thus difficult, so that this procedure may sometimes become unreliable, although vectorcardiogram (VCG) instead of ECG recordings are more suited and recommended to use for the R-peak detections when available (Mullinger & Bowtell, 2011).

Adapted from the context of EEG and also magnetoencephalography (MEG) source imaging, beamforming spatial filtering technique (Robinson and Vrba, 1999; Sekihara et al., 2001; van Drongelen et al., 1996; van Veen et al., 1997; van Veen and Buckley, 1988) is a well-known dipole scanning approach that also appears as a promising denoising technique, only recently considered in the context of EEG-fMRI studies (Brookes et al., 2009, 2008a; Mullinger and Bowtell, 2011). The beamforming technique is a spatial filtering technique, highly efficient when attenuating artefactual signals which have the different spatial origins from the underlying signal of interest, such as eye movements (Cheyne et al., 2006) and orthodontic metal braces (Cheyne et al., 2007) in MEG, and MRI related artefacts of EEG signals in EEG-fMRI (Brookes et al., 2008a), although the beamforming technique has been initially introduced for the source imaging technique in MEG and EEG studies (Robinson and Vrba, 1999; Sekihara et al., 2001; van Drongelen et al., 1996; van Veen et al., 1997; van Veen and Buckley, 1988). Specifically, the beamforming spatial filter rejects sources of signal variance that are not concordant in predetermined source locations in the brain based on the forward model. Consequently, it attenuates all unwanted source activities outside of the predetermined source location of interest in the data (e.g., eye movements) without having to specify the location or the configuration of these unwanted underlying source signals.

The residual artefacts from the GA and BCG artefacts are especially attenuated since their spatial topographies differ from the one corresponding to a dipolar current source located in the brain (Brookes et al., 2008a; Mullinger and Bowtell, 2011). The advantages of the beamforming denoising technique reside in the fact that it is a data-driven approach, which does not require to identify noise and signal components, and does not rely on the detection of ECG R-peaks. The beamforming denoising technique in EEG-fMRI (Brookes et al., 2008a; Mullinger and Bowtell, 2011) has initially been designed to minimize the residual GAs, and normally applied after correcting MRI related GA and BCG artefact as a postprocessing procedure. However, Brookes et al. (2008) tried the beamforming approach to correct the BCG artefacts without the conventional AAS BCG corrections after the GA corrections, and demonstrated that the beamforming technique indeed attenuated the BCG artefacts even though this previous work was preliminary and not extensive. Especially, in their previous study, this beamforming BCG denoising was examined in the EEG-fMRI data from two healthy adults when they were passively viewing 8.6Hz flashing checkerboard visual stimuli, and the performance of the beamforming denoising was evaluated by only successfully recovering the second harmonic signals (17.2Hz) in the source space in the primary visual cortex, when compared to the occipital sensor space signal. With the limited number of participants in the previous study, they did not extend their analysis to examine how much BCG artefacts were corrected by the beamforming BCG denoising technique, and did not investigate how much alpha event-related desynchronization was observed during the visual stimuli after the proposed beamforming BCG denoising.

In the present study, we proposed to systematically investigate the performance of this data-driven beamforming BCG denoising technique in EEG-fMRI. Therefore, the aim of this work was to investigate whether the BCG artefacts can be attenuated by only using beamforming techniques, i.e. without taking into account the ECG signal, and also to examine how this technique would preserve expected task-based induced brain signals in the source space while attenuating the BCG artefacts during an attentional network task (ANT). Furthermore, to clearly demonstrate the effects of the beamforming BCG artefact corrections on EEG task activities, we assessed the effects of beamforming on event-related desynchronization (ERD: reflecting a power decrease) in occipital alpha (8-13Hz) during the presentation of visual stimuli and motor beta (13-30Hz) during movement preparation before response onset during an attentional network task (ANT) (Fan et al., 2007, 2005). Specifically during the ANT task, the visual alpha ERD would be expected during the visual attention before the movement execution cue onset, whereas the motor beta ERD would be expected during the movement preparation/planning before the movement execution (Fan et al., 2007; Marshall et al., 2015), although the ANT task would also induce higher cognitive and attentional processing (alerting, orienting and executive control), which was beyond the scope of this paper. In this study, we analysed EEG-fMRI data in 20 healthy participants when they were performing the ANT task, and compared the four following BCG correction approaches: non-BCG corrected, AAS BCG corrected, beamforming+AAS BCG corrected, beamforming BCG corrected. We chose to investigate alpha ERD in the visual cortex and beta ERD in the motor cortex since the BCG artefacts typically obscure EEG signals below 20Hz (Bonmassar et al., 2002; Xia et al., 2014, 2013). Moreover, visual alpha ERD has been well documented during visual stimulations (Brookes et al., 2005; Scheeringa et al., 2011; Wilson et al., 2019), whereas reliable motor beta ERD has been reported during planning/preparation of the movement (Cheyne and Ferrari, 2013; Darvas et al., 2010; Hall et al., 2011; Muthukumaraswamy, 2010; Pfurtscheller et al., 1996; Pfurtscheller and Lopes da Silva, 1999; Takemi et al., 2013; Uji et al., 2018). We hypothesised that the beamforming BCG corrected data would recover visual alpha ERD and motor beta ERD to a similar extent, when compared to the conventional AAS BCG artefact corrections and beamforming+AAS BCG corrected data. Furthermore, we also hypothesised that both beamforming approaches would increase the signal to noise ratios (SNRs) in the source space when compared to those in the sensor space (Brookes et al., 2009; Hill et al., 2020; Sekihara et al., 2004), which should be another advantage of using beamforming techniques in EEG-fMRI.

## Methods

The EEG-fMRI data were acquired using a protocol previously published in Cross et al. (2021), and reused for our specific aims to investigate the performance of beamforming denoising techniques in simultaneous EEG-fMRI. The aim of the original study was to examine functional connectivity patterns across cognitive tasks of attention, vigilance, and working memory following sleep deprivation and a subsequent recovery nap. More details of the study designs and findings can be found in the previous article (Cross et al., 2021).

### Participants

20 young healthy participants (12 females, 8 males, age = 21.3 ± 2.5 years) took part in the study, and were screened for the absence of sleep disorders (insomnia, sleep apnea syndrome, hypersomnolence, restless legs syndrome, and parasomnias), neurological or psychiatric conditions (e.g. epilepsy, migraine, stroke, chronic pain, major depression, anxiety disorder, psychotic disorder) and current use of psychotropic medications or cannabis. All subjects provided written informed consent prior to the start of the study that was approved by the Central Research Ethics Committee of the Quebec Ministry of Health and Social Services. More details about the participants’ characteristics can be found in the previous study (Cross et al., 2021).

### Experimental procedures

Participants were invited for two different visits to the PERFORM Centre of Concordia University, Montréal, Canada. The two different visits were well-rested (WR) or sleep deprivation (SD) sessions. During the WR session, participants arrived at 7:00pm and were given a 9-hour sleep opportunity in a dark, quiet room, whereas during the SD session, they arrived at 8:00pm and stayed awake during the entire night in the laboratory. In both sessions, EEG-fMRI data acquisition began at 8:00am next day. In the WR session, the acquisition involved three different cognitive tasks (Attentional Network Task: Fan et al., 2002, 2005; Mackworth Clock Task: Lichstein et al., 2000; Loh et al., 2004; N-back task: Kirchner, 1958; Sweet, 2011). In the SD session, the first session involved the same three cognitive tasks, and then subjects were provided a 60-minute nap opportunity inside the MRI scanner. After 60 minutes, the participant was woken and during the final session (post recovery nap), they completed all the three cognitive tasks again. The EEG-fMRI data were acquired during three different cognitive tasks and one-hour nap (see Supplementary Figure S1A). The Attentional Network Task (ANT) is a task designed to test three different attentional networks: alerting, orienting, and executive control in higher cognitive and attentional processing (Fan et al., 2005, 2002). In this study, in order to investigate the performance of beamforming techniques in the EEG-fMRI, the ANT obtained from these three sessions was selected out of the three different cognitive tasks based on having clear baseline periods and clear event onsets of a visual stimuli and movement onset which would be predicted to induce alpha event related desynchronization (ERD) in the visual cortex and beta ERD in the motor cortex respectively (Fan et al., 2007; Marshall et al., 2015). Especially, we expected BCG artefacts to obscure these alpha and beta frequency band induced activities, since the main BCG artefact frequency profile is thought to be below 20Hz (Bonmassar et al., 2002; Xia et al., 2014, 2013). The ANT was believed to be the most suitable for the main aims of this paper when compared to the other two tasks.

### Attentional Network Task (ANT) paradigm

This task probes different attentional processes, such as alerting, orienting and executive control (Fan et al., 2007, 2005, 2002) (see Figure 1). The ANT consisted of a series of trials in which the participants were required to identify the direction (left or right) of the middle arrow in an array of five arrows within an upper or lower panel of the screen (Target cue). The middle arrow during the target cue was either congruent or incongruent to the direction of the other arrows. Different probing conditions (Probe cue) were displayed immediately prior to the appearance of the target cue, including both panels flashing (double cue; alerting), either panel flashing (valid or invalid spatial cues; orienting), or no cue preceding the target cue (no cue). One run of the ANT contained 145 trials in total consisting of 24 double cue trials, 73 valid cue and 24 invalid spatial cue trials, and 24 no cue trials. These trials were also divided into 73 congruent and 72 incongruent trials for the target cue. One run of the task lasted 13 minutes in duration, with each trial lasting around one second with a random jitter (range = 2-12 s, mean = 5 seconds) between each trial (see more details in Figure 1).

The ANT was run on a laptop computer using Inquisit software (Millisecond Software LLC, Seattle USA), displayed to the participant via a projector screen behind the MRI scanner. The participants responded via button presses made using a response pad attached to the fingers of the left hand. Behavioural outcomes of reaction time (ms) and accuracy (%; correct trials/number of trials) can be found in the supplementary Figure S1B.

**Figure 1.**
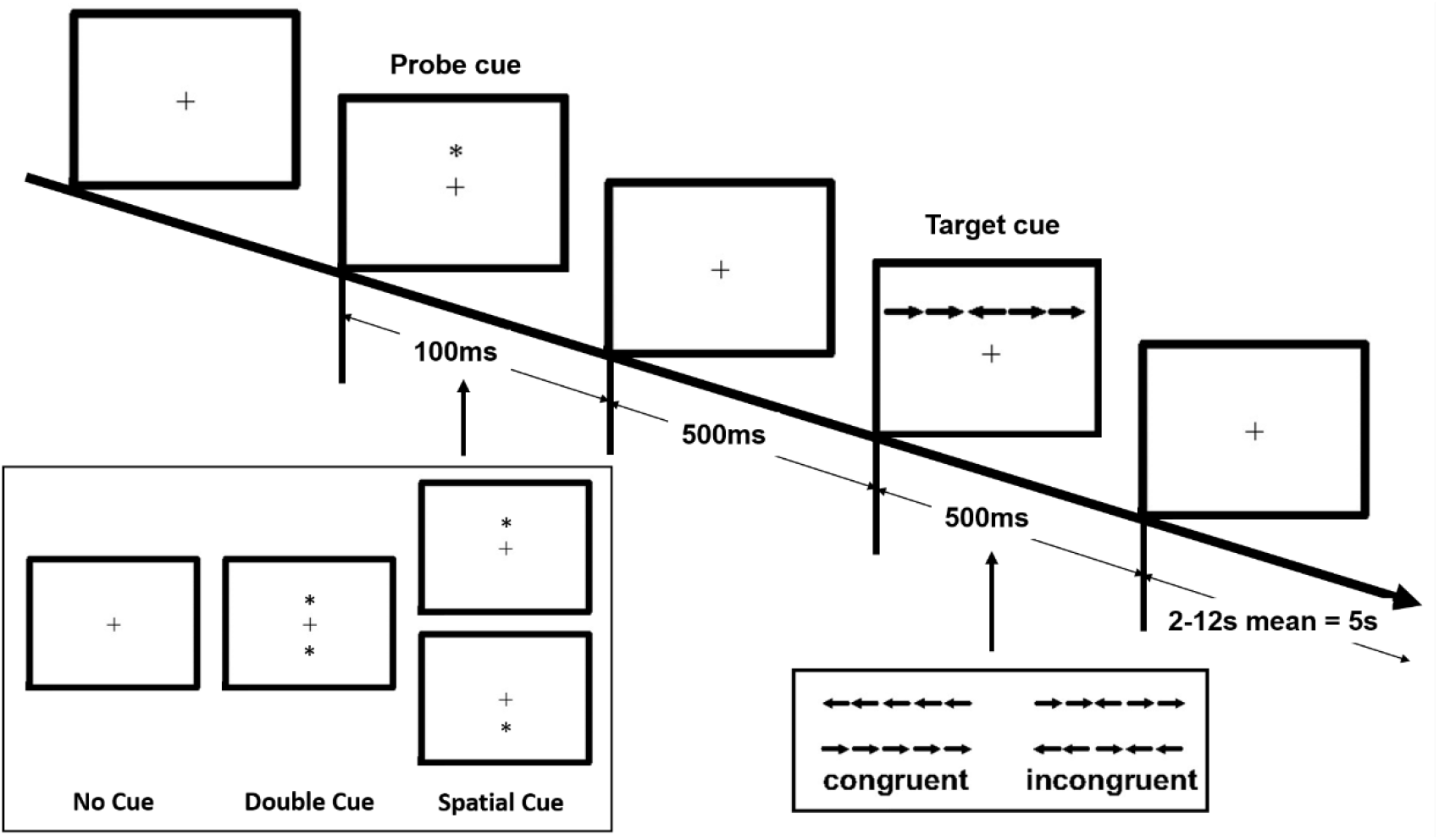
Schematic of attentional network task (ANT). In each trial, a fixation cross appears in the centre of the screen all the time. Depending on probe cue conditions, a probe cue (either no cue, double cue, or spatial cue) appears for 100ms. After a fixed duration (500ms), a target cue (the centre arrow) among other arrows (congruent or incongruent cue condition) is presented for 500ms at either an upper or lower panel of the screen, and participants are required to respond via button presses made using a response pad attached to the fingers of the left hand as quickly as possible, but no longer than 1900ms after the target cue onset. If participants respond with the correct direction of the middle arrow (left or right) within the time-window, the response is considered as a correct response. A post-target fixation period lasts for a variable duration (intertrial intervals from the offset of the target cue to the onset of the probe cue during the next trial are randomly jittered between 2000 and 12000ms, mean = 5s) between each trial. One run of the ANT contained 145 trials in total consisting of 24 double cue trials, 73 valid cue and 24 invalid spatial cue trials, and 24 no cue trials. These trials are also divided into 73 congruent and 72 incongruent trials for the target cue. One run of the ANT lasts around 13 minutes in duration.

### Simultaneous EEG-fMRI data acquisition

#### MRI data acquisition

MRI data were acquired with a 3T GE scanner (General Electric Medical Systems, Wisconsin, US) using an 8-channel head coil in the PERFORM Centre of Concordia University. Functional MRI were acquired using a gradient-echo echo-planar imaging (EPI) sequence (TR = 2500ms, TE = 26ms, FA = 90°, 41 transverse slices, 4-mm slice thickness with a 0% inter-slice gap, FOV = 192 x 192 mm, voxel size = 4 x 4 x 4mm^3^ and matrix size = 64 × 64). High-resolution T1-weighted anatomical MR images were acquired using a 3D BRAVO sequence (TR = 7908 ms, TI = 450 ms, TE = 3.06 ms, FA = 12°, 200 slices, voxel size = 1.0 × 1.0 × 1.0 mm, FOV = 256 × 256 mm). During all EEG-fMRI sessions, the Helium compression pumps used for cooling down MR components were switched off to reduce MR environment related artefacts infiltrating the EEG signal (Mullinger et al., 2008a; Rothlübbers et al., 2015). To minimize movement-related artefacts during the scanning, MRI-compatible foam cushions were used to fix the participant’ s head in the MRI head coil (Mullinger and Bowtell, 2011).

#### EEG data acquisition

EEG data were acquired using an MR compatible 256 high-density geodesic sensor EEG array (Electrical Geodesics Inc (EGI), Magstim-EGI, Oregon USA). The EEG cap included 256 sponge electrodes referenced to Cz that covered the entire scalp and part of the face. EEG data were recorded using a battery-powered MR-compatible 256-channel amplifier shielded from the MR environment that was placed next to the participant inside the scanning room. The impedance of the electrodes was initially maintained below 20kΩ and kept to a maximum of 70kΩ throughout the recording. Data were sampled at 1000Hz and transferred outside the scanner room through fibre-optic cables to a computer running the Netstation software (v5, EGI). The recording of EEG was phase-synchronized to the MR scanner clock (Sync Clock box, EGI), and all scanner repetition times (TRs) and participant responses were recorded in the EEG traces. Especially, the onset of every TR period was marked in the EEG data to facilitate MR gradient artefact correction. Electrocardiography (ECG) was also collected via two MR compatible electrodes placed between the 5th and 7th ribs and above the heart close to the sternum, and recorded at 1000Hz through a bipolar amplifier (Physiobox, EGI). After the EEG-fMRI data acquisition, EEG electrode locations were digitized using the EGI Geodesic Photogrammetry System (GPS) to facilitate individualized co-registration of electrode positions with each subject’s anatomical image (Russell et al., 2005).

### Data analysis

In this study, EEG-fMRI data during the ANT in 20 healthy participants were pre-processed, and compared with four different BCG correction approaches (non-BCG corrected, AAS BCG corrected, beamforming+AAS BCG corrected, beamforming BCG corrected) to examine how the beamforming techniques would attenuate BCG artefacts and preserve expected brain signals, notably visual alpha ERD and motor beta ERD, induced by the ANT task. The four different datasets consisting of non-BCG corrected and AAS BCG corrected sensor level data and beamforming source level datasets from either non-BCG corrected (beamforming BCG corrected) or AAS BCG corrected data (beamforming+AAS BCG corrected) were created in the manner described below.

For both EEG and ECG data, gradient artefacts (GA) were first corrected in BrainVision Analyzer2 (Brain Products Inc, Gilching Germany) with the standard AAS method, using sliding window templates formed from the averages of 21 TRs, which were subtracted from each occurrence of the respective artefacts for each electrode (Allen et al., 2000; Wilson et al., 2019). GA corrected data were subsequently bandpass filtered (EEG: 1-100 Hz with notch filter of 60Hz), and downsampled (500 Hz). Following the GA correction, cardiac R-peaks were automatically detected from the ECG recording in BrainVision Analyzer2, checked visually and corrected manually if necessary, since the ECG signal in the MRI scanner is distorted and has proven to be problematic (Chia et al., 2000; Mullinger et al., 2008b). These R-peak events were used to inform ballistocardiogram (BCG) correction of data recording inside the scanner.

From the same GA corrected datasets, two different datasets for each subject were created: non-BCG corrected and AAS BCG corrected data to investigate the effect of beamforming techniques on the BCG artefacts or remaining BCG artefacts after AAS (Brookes et al., 2008a). The AAS BCG artefact correction was performed using sliding template average-artefact subtraction (11 R-peak events per template) in BrainVision Analyzer2 (Mullinger et al., 2014).

For both non-BCG corrected and AAS BCG corrected data, two lowest rows of electrodes around the neck and face (69 electrodes) and ECG electrode were removed keeping 187 electrodes for further analysis. On the remaining 187 electrodes, noisy and bad electrodes, which were identified when the standard deviation of the normalized time-course of each electrode exceeded 5SD (Delorme and Makeig, 2004; Uji et al., 2019), were removed and interpolated using neighbour electrodes. Resulting data were then further downsampled into 250Hz and re-referenced to an average of all non-noisy channels, using EEGLAB software (https://sccn.ucsd.edu/eeglab) (Delorme and Makeig, 2004). After common average reference, all timepoints before and after the MRI scanning were removed, and further analyses were conducted on EEG data specifically during the MRI scanning.

#### EEG beamforming

To examine how the beamforming techniques would attenuate BCG artefacts and preserve expected brain signals, notably visual alpha ERD and motor beta ERD induced by the ANT task, we considered a linearly constrained minimum variance (LCMV) beamforming analyses (Brookes et al., 2008a; Uji et al., 2018; van Veen et al., 1997), using Fieldtrip toolbox implementation (http://www.ru.nl/neuroimaging/fieldtrip) (Oostenveld et al., 2011).

EEG electrode locations were first accurately digitized using the EGI Geodesic Photogrammetry System (GPS) to facilitate individualized co-registration of electrode positions with each subject’s anatomical image (Russell et al., 2005). The digitized EEG electrode positions were co-registered with the subjects’ T1-weighted anatomical image using fiducials and scalp surface fitting. A 4-layer (scalp, skull, cerebrospinal fluid (CSF), and brain), anatomically realistic volume conduction boundary element model (BEM) (Fuchs et al., 2002; Oostendorp and van Oosterom, 1991) was created by segmenting each subject’s T1-weighted images into skin, skull, CSF and brain compartments. In the 4-layer BEM head model, the electrical conductivity of the scalp, the skull, CSF and the brain was set to 0.33 S/m, 0.0042 S/m, 1 S/m and 0.33 S/m, respectively (Gramfort et al., 2010; Kybic et al., 2006). A template grid (5mm spacing) covering the whole brain volume was created from the MNI template brain (Colin27) and transformed to each subject’s anatomical space. The lead-field matrix at each location on the template grid in individual subject space was then calculated using their BEM, not constraining the orientation of the dipolar source placed on that point of the grid (Sekihara et al., 2004).

Using the LCMV beamforming spatial filtering, an estimate of the source amplitude vector 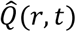 at source space location, r, and time, t, is made using a weighted sum of the electrical measurements recorded at each of the n electrodes such that:

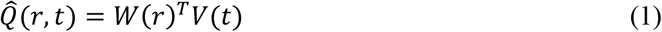

where V(t) is an n dimensional column vector of electric potential measurements made at time t, and W(r) is a 3×n matrix of weighting parameters that are tuned specifically to the location r.

Superscript T indicates a matrix transpose. The weighting parameters are calculated such that they minimise the total variance in the output signal 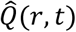, which will result in minimizing the influence of signals originated from any other location than the one of interest in r, whilst maintaining a constraint that any activity originating at the specific location r must remain in 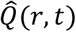.

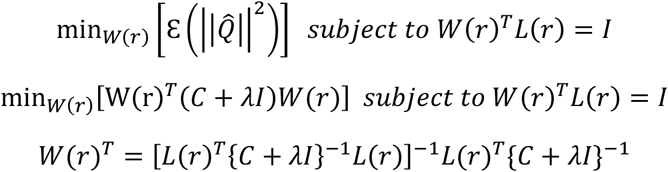

where L(r) represents a 3×n matrix in which the first, second and third columns of the lead field matrix for the position r, consisting of the electrical fields that would be measured at each of the n electrodes in response to a source of unit amplitude sited at location r and orientated in the x, y and z directions respectively. L(r) was therefore based on an EEG forward solution, (Sekihara et al., 2004; van Veen et al., 1997). ε denotes the mathematical expectation, whereas the matrix I represents the identity matrix. Here, *C* = ε(V(t)V(t)^T^) representing the n×n data covariance matrix calculated over a time-frequency window of interest using the entire epoched trials after removing noisy trials contaminated with large motion artefacts. λ is a regularization parameter and the regularized covariate matrix was calculated as (C + λl). By the reducing the regularization parameters it is possible to make the beamforming filter more spatially selective (Brookes et al., 2008b), and this would result in substantial suppression of the artefact (Litvak et al., 2010). Therefore, especially in the context of EEG-fMRI, a small regularization parameter of 0.01% is recommended to be used, therefore allowing more artefact suppressions (Brookes et al., 2008b; Litvak et al., 2010).

Both non-BCG corrected and AAS BCG corrected data were epoched from −2s to 3s relative to the probe cue events for identifying the alpha ERD VE location in the visual region (visual epoched data), and epoched from −4s to 2s relative to the movement onsets for the beta ERD VE location in the motor region (movement epoched data). To achieve cleaner EEG data, around 5-10% trial rejection rate is usually recommended (Delorme et al., 2007), and by applying amplitude threshold of ±500 μV on the AAS BCG corrected data, noisy EEG trials, that were contaminated with large motion artefacts, were detected and removed (Delorme et al., 2007; Uji et al., 2019). This resulted in a group mean (±SD) number of remaining trials of 132 ± 7 trials for each run for the visual epoched trials and 117 ± 15 trials for each run for the movement epoched trials for further analysis. The same numbers of trials were used for both AAS BCG corrected and non-BCG corrected data analysis.

In order to improve signal to noise ratios (SNRs) of the beamforming spatial filtering, an optimum source orientation/direction needs to be calculated instead of using 3D vector (x, y, z) at the source space location r (Sekihara et al., 2004). The optimal orientation opt(r) at the position r can be computed by the eigenvector corresponding to the minimum generalized eigenvalue of the matrix [L(r)^T^(C + λl)^-2^L(r)] with the metric [L(r)^T^(C + λI)^-1^L(r)] (Sekihara et al., 2004; Sekihara and Nagarajan, 2008). Once opt(r) is obtained, the beamforming weight vector (1D vector) can be calculated.

A scalar LCMV beamforming (Sekihara et al., 2004; Sekihara and Nagarajan, 2008) with individual BEM head models using the regularization parameter of 0.01% (Brookes et al., 2008b; Litvak et al., 2010) was carried out for each subject to calculate beamforming weights (weights of each EEG channel at each lead-field virtual electrode (VE) position r in the brain) derived from the entire broadband data (1-100Hz) and a covariance matrix calculated using the entire clean dataset of either visual or movement epoched trials after removing noisy trials (Brookes et al., 2008a). For every VE location within the source grid space, the estimated VE trial time-courses were then bandpass filtered using 6th-order Butterworth filters in the frequency band of interest (Alpha: 8-13Hz; Beta: 13-30Hz).

To identify the VE locations of interest, related to the induced task neural activity, i.e. visual alpha ERD and motor beta ERD, a pseudo-T statistic can be calculated following:

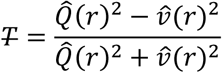

where 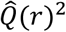 represents the Beamformer power estimated in r during an active time window, whereas 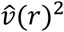 represents the Beamformer power estimated in r during a control baseline time window (Brookes et al., 2008a; Hillebrand and Barnes, 2005; Robinson and Vrba, 1999).

Specifically, to detect the alpha ERD VE location following visual stimuli, we calculated alpha power during the active time window of 0.1s to 1.1s after the probe cues, and the baseline period was defined between −1.5s to −0.5s prior to the probe cues. We then identified the minimum peak of the pseudo-T statistic map reflecting the largest alpha band power decrease, as the alpha VE location for each subject. To detect the beta ERD VE location during movement preparation/planning, we calculated beta power during the active time window between −1.25 s to −0.25 s prior to the movement onsets, whereas the baseline time window was defined between −3.0s to −2.0s prior to the movement onsets. We then identified the minimum peak of the pseudo-T map reflecting the largest beta band power decrease, as the beta VE location in the brain for each subject. From these pseudo-T maps, VE locations were separately identified for each subject. The alpha ERD VE location was indeed detected in the visual cortex (visual alpha VE) from the visual epoched data, whereas the beta ERD VE location was identified in the motor cortex (motor beta VE) from the movement epoched data.

A broadband (1-100 Hz) time-course of neural activity during the whole MRI scanning period was then also calculated from these two subject-specific VE locations, using equation (1). The beamforming weights of the visual alpha VE and motor beta VE were applied to the entire time-course of the whole 187 scalp electrodes during MRI scanning for R-peak locked analysis and ANT task-based analysis.

For further event-locked analysis, namely R-peak locked analysis and then ANT task-based analysis, we considered four data sets (non-BCG corrected; AAS BCG corrected; beamforming BCG corrected; beamforming+AAS BCG corrected) consisting of non-BCG corrected and AAS BCG corrected sensor level data and broadband (1-100 Hz) source level time-courses in the visual and motor beamforming VEs estimated from either non-BCG corrected or AAS BCG corrected data.

#### R-peak locked analysis

##### Sensor-level analysis

To investigate the effects of BCG artefacts on the EEG data at the sensor level, both non-BCG corrected and AAS BCG corrected data were epoched from −5s to 5s relative to every detected R-peak event (Group mean number (± SD) of R-peak events for each run = 870 ± 126 events). Two main analyses were conducted: (i) R-peak locked time-course analysis and (ii) R-peak locked time-frequency analysis.

For the R-peak locked time-course analysis, each R-peak locked time-course for each electrode was extracted, and each 10s epoch was demeaned by subtracting the mean value of the 10s epoch and divided by the standard deviation of the 10s epoch for normalized z-score transformation, for each selected epoch. This z-score transformation allowed us to make comparisons in the same scale since there would be differences in the amplitudes and signal to noise ratios (SNR) in the sensor and source level signals. Then calculated z scores were averaged across epochs first and then across subjects, for each electrode in both non-BCG corrected and AAS BCG corrected data.

In the R-peak locked time-frequency analysis, time-frequency spectrograms of each 10s epoch for each channel were calculated using a multitaper approach (Scheeringa et al., 2011; Uji et al., 2018) using the Fieldtrip toolbox. Windows of 0.8s duration were moved across the data in steps of 50ms, resulting in a frequency resolution of 1.25 Hz, and the use of three tapers resulted in a spectral smoothing of ±2.5Hz. The calculated time-frequency representations were then transformed in the scale of dB by 10*log10, then demeaned by subtracting the mean power in each frequency during the 10s epoch to calculate power fluctuations around a single R-peak, and averaged across epochs first and then across subjects for each electrode in both non-BCG corrected and AAS BCG corrected data.

##### Source-level analysis

The same R-peak locked time-course analysis and time-frequency analysis was conducted on the beamforming estimated time-course signals from the AAS BCG corrected (beamforming+AAS BCG corrected) and non-BCG corrected (beamforming BCG corrected) data. The whole time-course signals during the MRI scanning were estimated from the two VE locations using equation (1). We then epoched the whole visual VE and motor VE signals from −5s to 5s relative to every R-peak event from the AAS BCG corrected data. Both R-peak locked time-courses and time-frequency representations were averaged across epochs and then across subjects for both VE locations estimated from both the AAS BCG corrected (beamforming+AAS BCG corrected) and non-BCG corrected data (beamforming BCG corrected).

#### ANT task-based analysis

To clearly demonstrate the effects of BCG artefacts correction on resulting EEG signals of interest, we decided to investigate visual alpha event-related desynchronization (ERD) in response to the visual target cues and motor beta ERD during movement preparation/planning before the movement onset, instead of examining higher cognitive and attentional processing (alerting, orienting and executive control) expected during an ANT task (Fan et al., 2007, 2005).

The time-frequency analysis was chosen to investigate the effects of the BCG artefacts in the time-frequency domains, which normally obscure the EEG signals below 20Hz. This analysis was conducted on both visual and movement epoched data separately for both non-BCG corrected and AAS BCG corrected data. In the sensor space, using the same multitaper approach described above, we calculated the time-frequency representations in the visual epoched data for each channel, and the time-frequency spectrograms were converted to display power change in the task induced neural activity relative to baseline in the scale of dB (10*log10(signal/baseline)) with the baseline period between −1.5s to −0.5s prior to the probe cues. For the movement epoched data, the same analysis was conducted using the baseline period between −3.0s to −2.0s prior to the movement onsets.

In the source-space analysis, both visual and movement epoched trial time-courses were estimated from the two VE locations using equation (1) with respective beamforming spatial filter and epoched EEG signals from the AAS BCG corrected and non-BCG corrected data. The same time-frequency analysis was conducted for the visual alpha VE signals using the baseline period between −1.5s to −0.5s prior to the probe cues, whereas the motor beta VE signals were analysed using the baseline period between −3.0s to −2.0s prior to the movement onsets in the same time-frequency analysis.

#### Statistical analysis

Non-parametric cluster-based permutation tests (Fieldtrip) (Maris, 2012; Maris and Oostenveld, 2007; Nichols and Holmes, 2002) were used to examine significant differences comparing the TFRs in the sensor and source space from the R-peak locked and ANT task-based analysis. To do so, when comparing two BCG corrections strategies among the four proposed investigations (e.g. beamforming+AAS BCG corrected versus AAS BCG corrected), we considered all normalized TFR maps from all 20 subjects to build a time-frequency t-map of the difference between the two approaches, therefore taking into account inter-subject variability. For each combination of methods to be tested, statistical inference was based on a non-parametric cluster-based permutation test, which resulted in time-frequency t-map clusters obtained when first applying a threshold of alpha level of 0.001. From this thresholded time frequency t-map, the t-values were summed per cluster and then used as the test statistic. A randomization distribution of this test statistic was determined by randomly exchanging 1000 times, through permuting the two conditions among the N=20 subjects and the permutation p-value of the cluster of interest was approximated by a Monte Carlo estimate. This was done by building a null distribution considering all possible permutations given the number of subjects (N=20), and for each randomization only the maximal test statistic was retained. An observed cluster was deemed significant if it fell outside the central 95% of this randomization null distribution, corresponding to a two-sided random effect test with 5% false positives, corrected for the multiple comparisons across times and frequencies. The significant spectral-temporal cluster masked the raw time-frequency representation difference. The permutation test has advantages over the Bonferroni correction for multiple comparisons because the Bonferroni correction assumes that all measures are independent, an assumption that is too strong and weakens the power of the statistical test. In contrast, the permutation test considers the true dependency among all of the measures.

#### Signal to noise ratio (SNR) comparisons during ANT

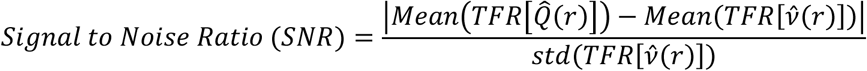

Where 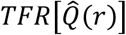 denotes the time-frequency power at either EEG electrode or VE location of interest, within the frequency range of interest during the active time window, whereas 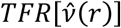 represents the time-frequency power at either EEG electrode or VE location of interest, within the frequency range of interest during the control baseline time window. *Mean* and *std* denotes respectively the mean and standard deviation computed over all coefficients of the selected time-frequency area.

Since the alpha and beta ERD exhibits power reduction when compared to the baseline, signal to noise ratios (SNRs) at the sensor and source levels were calculated as the absolute value of the difference. Then group mean SNR value was calculated by averaging SNR value across all trials first and then across subjects as suggested in Hill et al. (2020), for each electrode (non-BCG corrected; AAS BCG corrected) and each VE location (beamforming+AAS BCG corrected; beamforming BCG corrected). The active and control baseline time windows were consistent throughout the data analysis. For the visual alpha (8-13Hz) ERD, the active window was the time period between 0.1s to 1.1s as compared with the baseline window of −1.5s to −0.5 s relative to the probe cue onset, whereas for the motor beta (13-30Hz) ERD, the active window was the time period between −1.25s to −0.25s as compared with the baseline window of −3s to −2s relative to the movement onset.

### Data and code availability

All data used in this study will be available via a request to the corresponding author (MU). Open access software used in this study is available at https://sccn.ucsd.edu/eeglab (EEGLAB) & http://www.ru.nl/neuroimaging/fieldtrip (FieldTrip). Custom analysis code used in this study will be available via a request to the corresponding author (MU).

## Results

### Ballistocardiogram (BCG) artefact attenuation

#### BCG artefact corrections in the sensor space

The group mean R-peak locked time-courses from the non-BCG corrected data were represented on 17 electrodes selected from the EGI geodesic electrodes, which have the equivalent locations from the international 10-20 EEG system (see Figure 2). The group mean R-peak locked time-courses from the AAS BCG corrected data on the same 17 electrodes, are represented in Figure 3. The fluctuations of the non-BCG artefact time-courses resulted in z-scores values ranging between 0.5 to 1 at the time of R-peaks (Figure 2). When considering the resulting residual BCG artefact after applying the standard AAS correction method, we obtained peak z-score values ranging between 0.05 to 0.1 (Figure 3), demonstrating that the conventional AAS approach enabled around 10-fold improvement in suppression of BCG artefacts.

**Figure 2.**
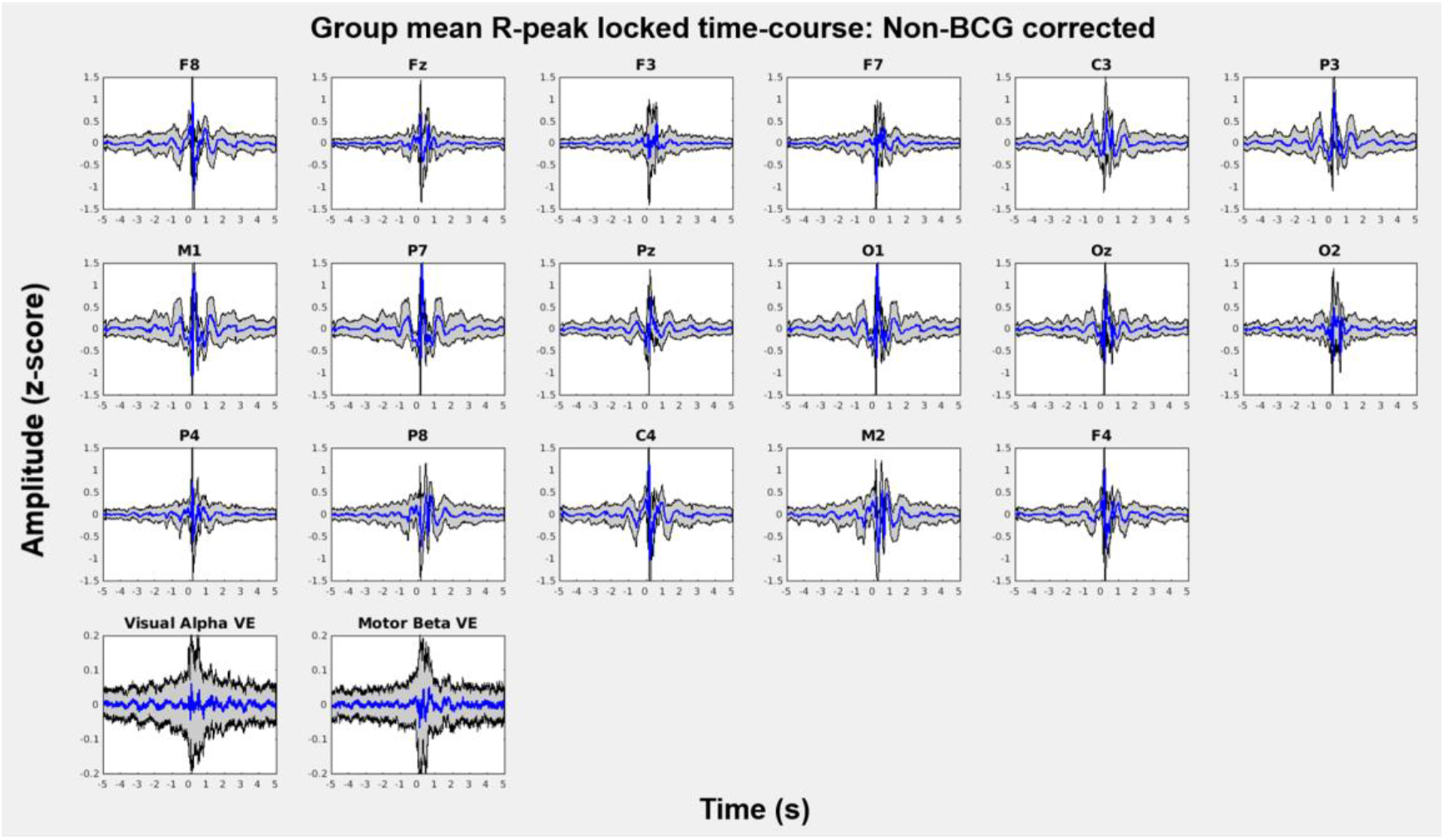
Group mean ± SD (N = 20 participants) z-scores of ballistocardiogram (BCG) artefact related time–courses time-locked at individual R-peak events from ECG recording during each run (3 runs of 13 min in total). The time-courses were measured from no BCG corrected data on 17 electrodes selected from an EGI geodesic 256 EEG system and 2 virtual electrodes (VEs) in the visual and motor cortex after applying beamforming techniques on the non-BCG corrected data (beamforming BCG corrected). Specifically, E2 (F8), E21 (Fz), E36 (F3), E47 (F7), E59 (C3), E87 (P3), E94 (M1), E96 (P7), E101 (Pz), E116 (O1), E126 (Oz), E150 (O2), E153 (P4), E170 (P8), E183 (C4), E190 (M2), and E224 (F4) electrodes have the equivalent locations in an international 10-20 EEG system from the EGI geodesic 256 EEG system. It needs to be noted that the scales of y-axis at the sensor level are within ±1.5 z-scores, whereas those at the source level are within ±0.2 z-scores.

**Figure 3.**
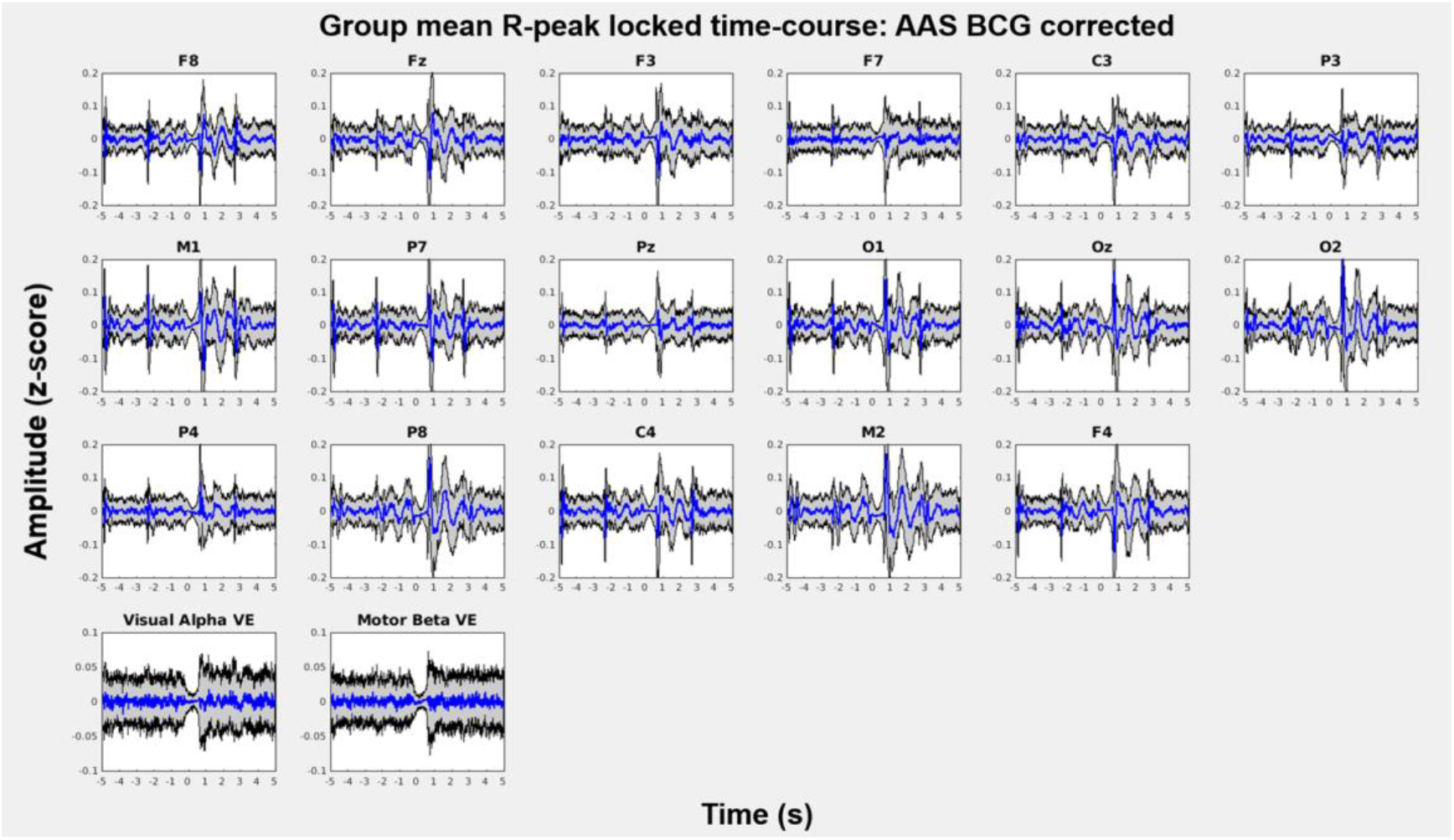
Group mean ± SD (N = 20 participants) z-scores of ballistocardiogram (BCG) artefact corrected time-courses time-locked at individual R-peak events from ECG recording during each run (3 runs in total). The time-courses were measured from the BCG corrected data (AAS approach) on 17 electrodes selected from an EGI geodesic 256 EEG system and 2 virtual electrodes (VEs) in the visual and motor cortex after applying beamforming on the AAS BCG corrected data (beamforming+AAS BCG corrected). Specifically, E2 (F8), E21 (Fz), E36 (F3), E47 (F7), E59 (C3), E87 (P3), E94 (M1), E96 (P7), E101 (Pz), E116 (O1), E126 (Oz), E150 (O2), E153 (P4), E170 (P8), E183 (C4), E190 (M2), and E224 (F4) electrodes have the equivalent locations in an international 10-20 EEG system from EGI geodesic 256 EEG system. It needs to be noted that after the AAS BCG corrections, the scales of y-axis at the sensor level are within ±0.2 z-scores, whereas those at the source level are within ±0.1 z-scores.

The group mean time-frequency representations (TFRs) time-locked at the individual R-peak events, when applied on non-BCG corrected data, are presented in Figure 4 on the selected 17 electrodes. The group mean TFRs time-locked at the individual R-peak events from the AAS BCG corrected data, are presented in Figure 5. When considering non-BCG corrected data, the power fluctuations of the BCG artefact were observed at the peak power of around 1 dB, especially below 20Hz, at the time of R-peaks (Figure 4). After applying the standard AAS BCG correction, these power fluctuations were reduced to the peak power of around 0.2 dB at the time of R-peaks especially below 20Hz (Figure 5). In Figure 6, the cluster-based permutation tests revealed significant differences (p < .05) between the AAS BCG corrected and non-BCG corrected data for all the selected 17 electrodes, confirming that the standard AAS technique removed the BCG artefacts significantly. Although some of the residual artefacts can be observed above 20Hz, the AAS method, in general, demonstrated overall good suppression of BCG artefacts in time-frequency domains by comparing the TFRs of the non-BCG corrected and AAS BCG corrected data.

**Figure 4.**
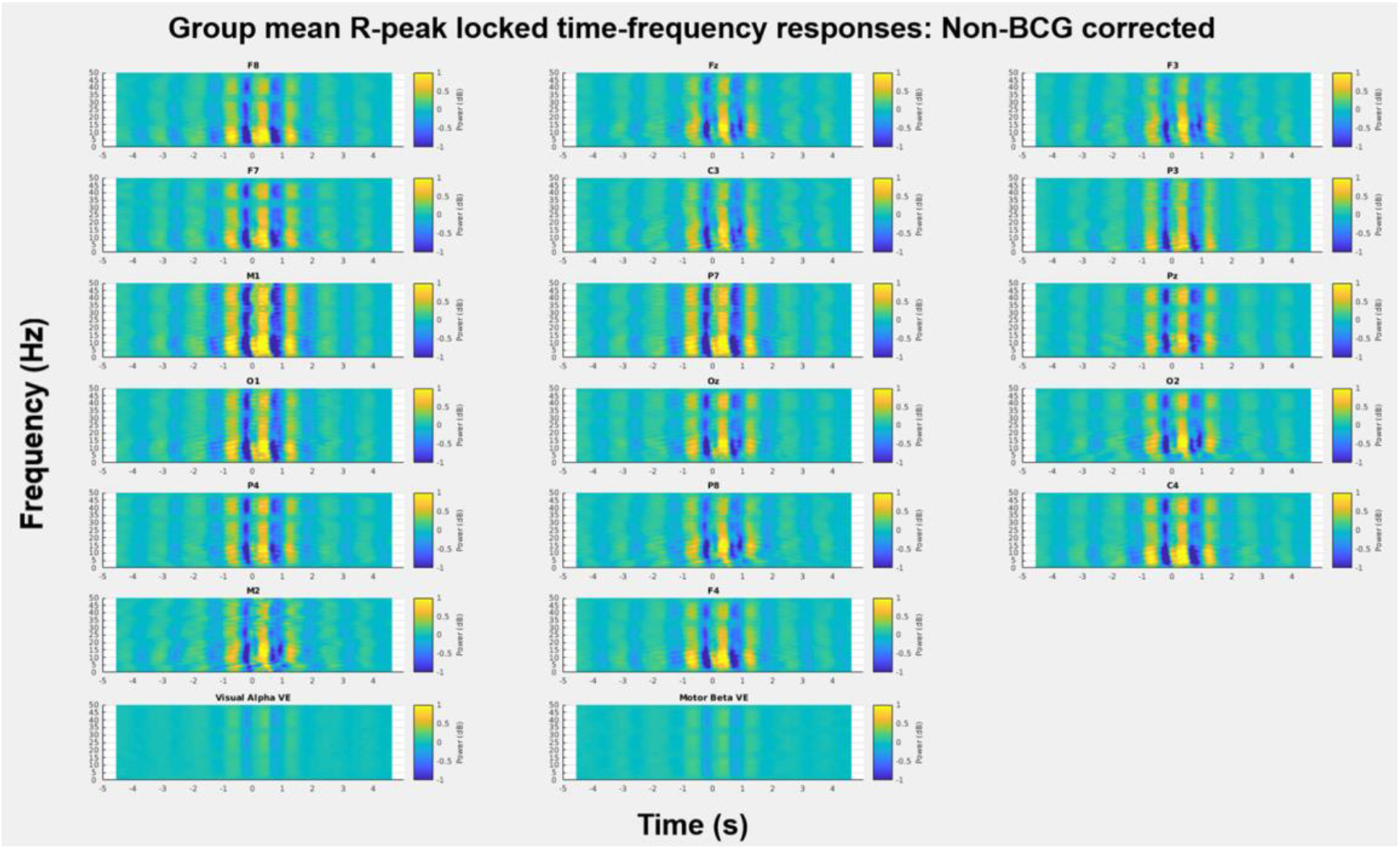
Group mean (N = 20) time-frequency representations (TFRs) of ballistocardiogram (BCG) artefact related signals time-locked at individual R-peak events from ECG recording during each run (3 runs in total). The TFRs were calculated from non-BCG corrected data on selected 17 electrodes from an EGI geodesic 256 EEG system and 2 virtual electrodes (VEs) in the visual and motor cortex after applying beamforming techniques on the non-BCG corrected data (beamforming BCG corrected). Specifically, E2 (F8), E21 (Fz), E36 (F3), E47 (F7), E59 (C3), E87 (P3), E94 (M1), E96 (P7), E101 (Pz), E116 (O1), E126 (Oz), E150 (O2), E153 (P4), E170 (P8), E183 (C4), E190 (M2), and E224 (F4) electrodes have the equivalent locations in an international 10-20 EEG system from EGI geodesic 256 EEG system. It needs to be noted that the scales of colour bars in this figure are within ±1 dB.

**Figure 5.**
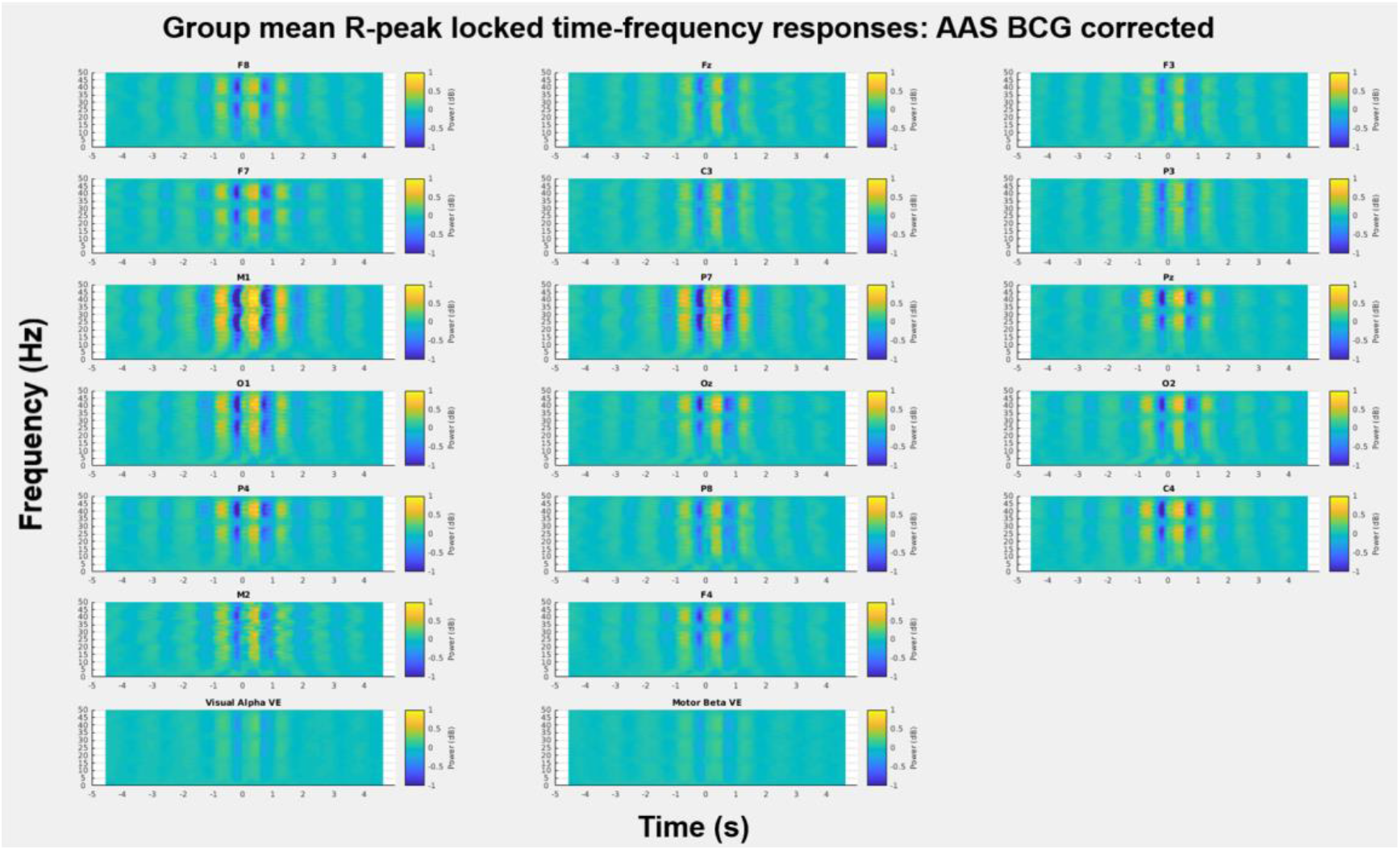
Group mean (N = 20) time-frequency representations (TFRs) of ballistocardiogram (BCG) artefact corrected signals time-locked at individual R-peak events from ECG recording during each run (3 runs in total). The TFRs were calculated from the BCG corrected data (AAS approach) on selected 17 electrodes from an EGI geodesic 256 EEG system and 2 virtual electrodes (VEs) in the visual and motor cortex after applying beamforming techniques on the AAS BCG corrected data (beamforming+AAS BCG corrected). Specifically, E2 (F8), E21 (Fz), E36 (F3), E47 (F7), E59 (C3), E87 (P3), E94 (M1), E96 (P7), E101 (Pz), E116 (O1), E126 (Oz), E150 (O2), E153 (P4), E170 (P8), E183 (C4), E190 (M2), and E224 (F4) electrodes have the equivalent locations in an international 10-20 EEG system from EGI geodesic 256 EEG system. It needs to be noted that the scales of colour bars in this figure are within ±1 dB.

**Figure 6.**
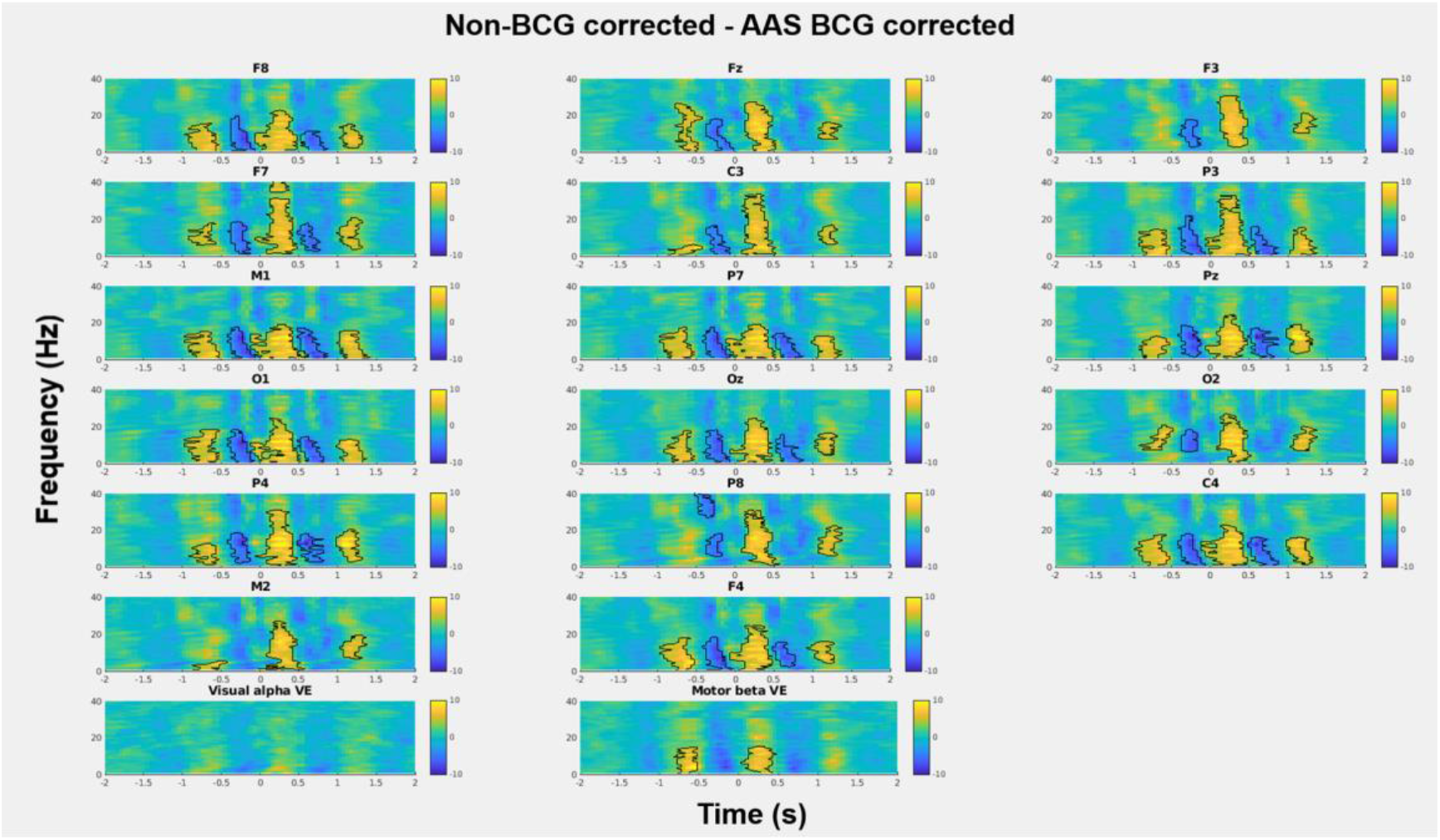
The t-maps of differences in the time-frequency representations (TFRs) between the AAS BCG corrected and non-BCG corrected data in both sensor and source levels (Visual and Motor VEs). Figures demonstrate t-values and black contour representing the significant clusters from cluster-based permutation tests (p < .05). Yellow denotes a positive and blue denotes a negative t-values, suggesting a significant reduction of the BCG artefacts when applying AAS, as opposed to non-BCG correction, and overall significant improvement of AAS methods for signals below 20Hz.

#### Beamforming approach

Beamforming spatial filtering was applied to both AAS BCG corrected (beamforming+AAS corrected) and non-BCG corrected (beamforming BCG corrected) data, in order to investigate how this technique would attenuate BCG artefacts. The virtual electrode (VE) locations were determined by the analysis of the ANT task results which consisting in the identification of one VE in the visual cortex (Visual Alpha VE) and one VE in the motor cortex (Motor Beta VE) exhibiting the largest ERD results. A specific VE location (Visual and Motor) was found for every participant, using either beamforming+AAS corrected data or beamforming BCG corrected, and converted into MNI coordinates. The MNI coordinates of the group mean visual alpha VE location (±SE) was [−25±4, −75±1, −8±3] mm [MNI:x,y,z] for the AAS BCG corrected data and [2±4, −96±1, 9±3] mm [MNI:x,y,z] for the non-BCG corrected data. The MNI coordinates of the group mean motor beta VE location (±SE) was [45±5, −44±3, 54±2] mm [MNI:x,y,z] for the AAS BCG corrected data and [55±7, −35±3, 50±2] mm [MNI:x,y,z] for the non-BCG corrected data (Figure 7, see details below in Attentional Network Task (ANT) task activity).

**Figure 7.**
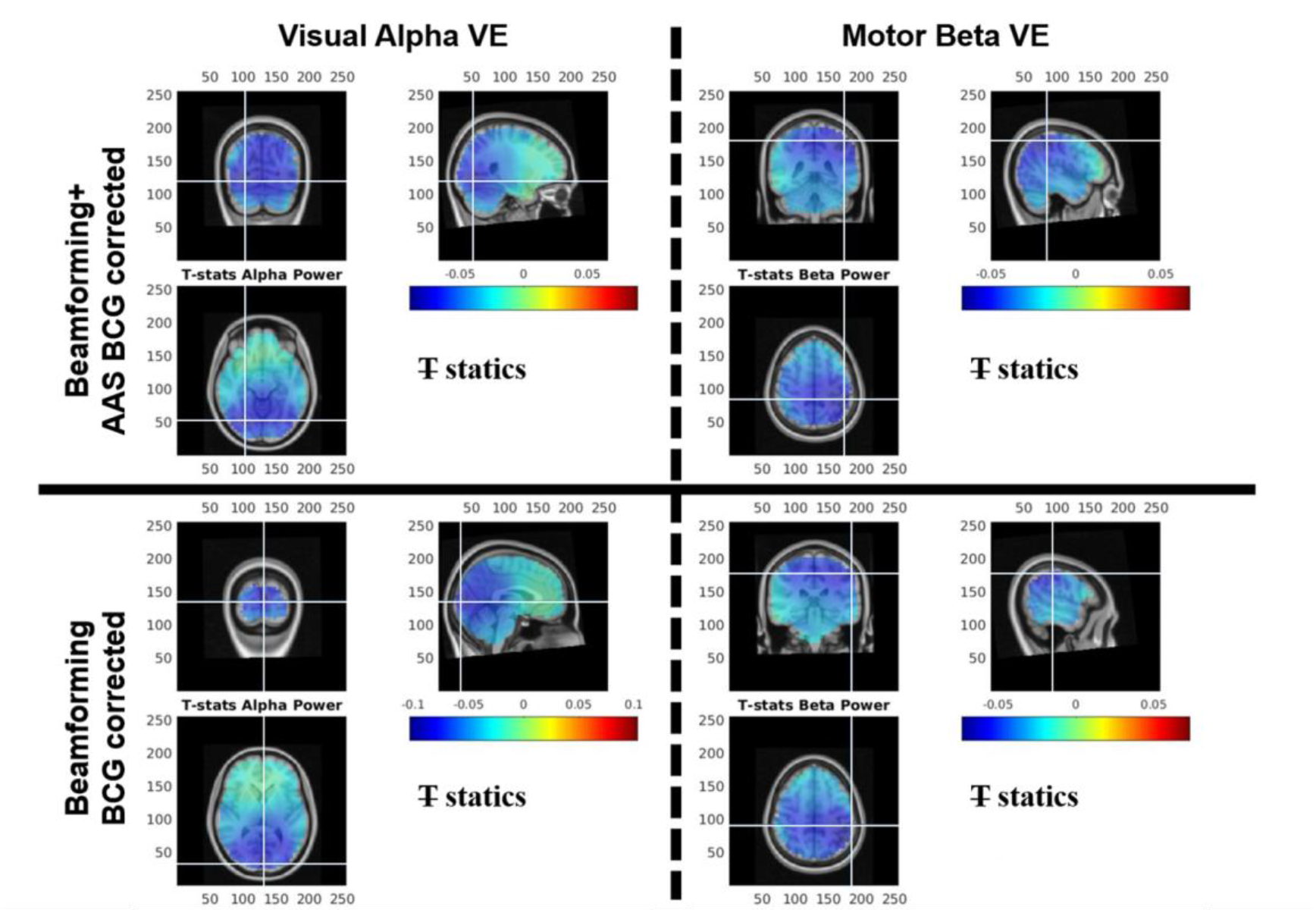
Group average (N = 20) pseudo-T statistic beamforming maps showing (Left column) regions exhibiting power decreases in alpha frequency (8-13Hz) during the active window (0.1-1.1s) as compared with the baseline window (−1.5 to −0.5 s) relative to the probe cue onset from the AAS BCG corrected (Top) and non-BCG corrected data (Bottom), whereas (Right column) regions exhibiting power decrease in beta frequency (13-30Hz) during the active window (−1.25 to −0.25s) as compared with the baseline window (−3 to −2 s) relative to the movement onset from the AAS BCG corrected (Top) and non-BCG corrected data (Bottom). The crosshairs represent the group average of the individual VE locations found in visual cortex for the alpha frequency activity (left column) and in the motor cortex for the beta frequency activity (right column). The group mean visual alpha VE locations across subjects (±SE) was found at [−25±4, −75±1, −8±3] mm [MNI:x,y,z] for the beamforming+AAS BCG corrected data and [2±4, −96±1, 9±3] mm [MNI:x,y,z] for the beamforming BCG corrected data (see crosshairs), whereas the group mean motor beta VE locations (±SE) was found at [45±5, −44±3, 54±2] mm [MNI:x,y,z] for the beamforming+AAS BCG corrected data and [55±7, −35±3, 50±2] mm [MNI:x,y,z] for the beamforming BCG corrected data, both of which were contralateral to the left hand button presses (see crosshairs).

Using those subject specific locations, the group mean VE time-courses time-locked at the individual R-peak events were extracted from the non-BCG corrected data, whose original data clearly demonstrated the presence of large BCG artefacts at the sensor level (see Figure 2). The group mean VE time-courses time-locked at the individual R-peak events were also calculated from the AAS BCG artefact corrected data, as illustrated in Figure 3. It is important to mention that the fluctuations of the remaining BCG artefact on the VE time-courses from the beamforming BCG corrected data, resulted in the peak z-score values around 0.05 (Figure 8), when compared to the original BCG artefacts which had the peak z-scores between 0.5 to 1 at the R-peak events in the sensor space of the non-BCG corrected data. Moreover, the fluctuations of the remaining BCG artefact from the beamforming BCG corrected data was similar to those of the beamforming+AAS BCG corrected data showing the peak z-score values around 0.05 (Figure 8), suggesting that whether AAS was considered or not at the sensor level, spatial filtering using beamforming was able to reach the similar amount of BCG artefact removal.

**Figure 8.**
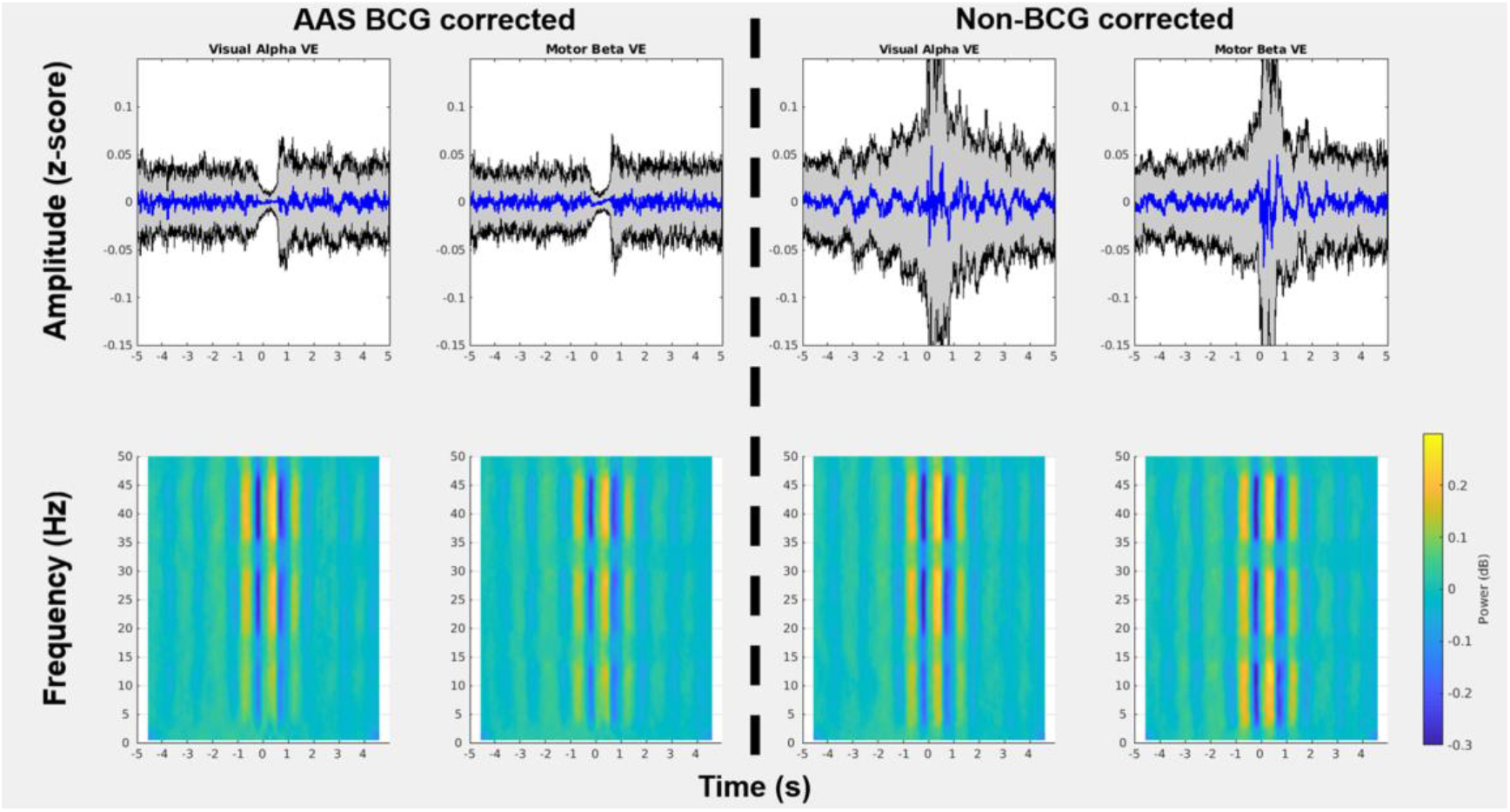
Group mean (N = 20) time-courses (±SD) and time-frequency representations (TFRs) of ballistocardiogram (BCG) artefact related signals time-locked at individual R-peak events from 2 virtual electrodes (VEs) in the visual and motor cortex after applying beamforming techniques on either AAS BCG corrected (beamforming+AAS BCG corrected) or non-BCG corrected data (beamforming BCG corrected). It needs to be noted that the scales of the y-axis in the time-course amplitude are within ±0.15 z-scores, whereas those of the colour bars in the TFR are within ±0.3 dB.

The group mean VE TFRs time-locked at the individual R-peak events were extracted from the beamforming BCG corrected data, whose original data clearly demonstrated remaining BCG artefacts at the sensor level (see Figure 4), whereas the group mean VE TFRs time-locked at the individual R-peak events were also calculated from the beamforming+AAS BCG artefact corrected data in Figure 5. The power fluctuations of the remaining BCG artefact from the VE signals of the non-BCG corrected data were observed at the peak power of around 0.2 dB at the time of R-peaks (Figure 8), resulting in a large attenuation of the BCG artefacts which were originally exhibiting a peak power of around 1 dB at the R-peak events, especially below 20Hz. In Figure 9b, the cluster-based permutation tests revealed significant reduction (p < .05) of the BCG artefact between the beamforming BCG corrected data (visual alpha VE, motor beta VE) and corresponding non-BCG corrected electrodes (Oz, C4), confirming that the beamforming technique significantly removed the BCG artefacts, even from non-corrected data.

**Figure 9.**
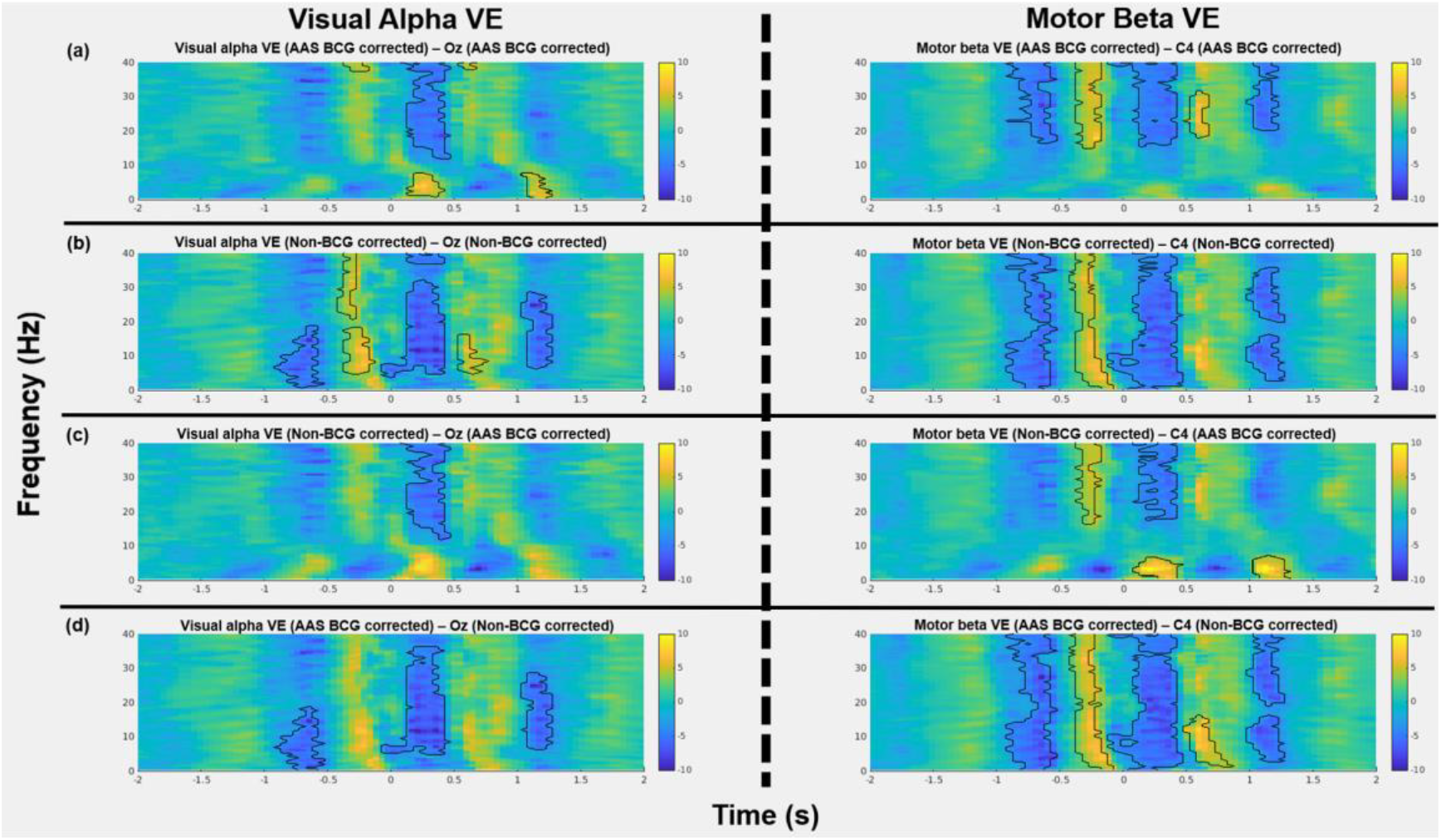
R-peak locked analysis: The t-maps of differences between the time-frequency representations (TFRs) for various comparisons, (a) between beamforming+AAS BCG corrected and corresponding AAS BCG corrected EEG electrodes (Oz for visual, C4 for motor), (b) between beamforming BCG corrected and corresponding non-BCG corrected EEG electrodes, (c) between beamforming BCG corrected and corresponding AAS BCG corrected EEG electrodes, (d) between beamforming+AAS BCG corrected and corresponding non-BCG corrected EEG electrodes, in the visual alpha VE in the left column and the motor beta VE in the right column. Figures demonstrate t-values and black contour representing the significant clusters from cluster-based permutation tests (p < .05). Yellow denotes a positive and blue denotes a negative t-values.

By comparing the standard AAS approach (AAS BCG corrected data) and beamforming+AAS BCG corrected data (Figure 9a), the cluster-based permutation tests revealed significant differences (p < .05) between the beamforming+AAS BCG corrected data (visual alpha VE, motor beta VE) and corresponding AAS BCG corrected electrodes (Oz, C4), suggesting the beamforming spatial filtering could also further improve the BCG artefact suppressions even after the standard AAS approach in the sensor space. Furthermore, the cluster-based permutation tests (Figure 9c) revealed significant differences (p < .05) between the beamforming BCG corrected data (visual alpha VE, motor beta VE) and corresponding AAS BCG corrected electrodes (Oz, C4), demonstrating that the performance of this beamforming BCG denoising for the non-BCG corrected data (Figure 8) was significantly greater than the level of the conventional AAS approach (Figure 5). This finding indicates that the proposed data-driven beamforming denoising approach allows significantly attenuating the BCG artefacts without relying on R-peak detections from ECG recording. Lastly, by comparing the performance of both beamforming approaches (beamforming+AAS BCG corrected, beamforming BCG corrected), the permutation tests revealed that the beamforming+AAS BCG corrected data contained significantly less BCG artefacts than the beamforming BCG corrected data in the motor beta VE (p < .05), although there was no significant difference between them in the visual alpha VE, indicating some advantages of applying the beamforming technique on the corrected data, when compared to solely applying beamforming on the uncorrected data (Figure 6).

### Attentional Network Task (ANT) task activity

#### Visual alpha event-related desynchronization (ERD)

##### Analysis at the sensor level

Figure 10 shows the group mean TFRs measured in selected occipital and parietal electrodes, reporting alpha (8-13Hz) power change in response to the target cues which occurred at 600ms after the probe cue onset (either no cue, double cue, or spatial cue). Results are reporting when considering results obtained for the AAS BCG corrected data (top row) and non-BCG corrected data (bottom row). Despite of the large broadband increases in power (red vertical stripes occurred after around 1s in response to the probe cue onset) demonstrated the motion artefacts caused by the button press for the task responses (see Supplementary Figure: S2), alpha power decrease (alpha ERD) was still observed within the time window before 1s when compared to baseline time-window (−1.5s to −0.5s prior to the probe cue onset) in the AAS BCG corrected data (see top row in Figure 10). However, this alpha ERD was difficult to identify in the same electrodes from the non-BCG corrected data (see bottom row in Figure 10) as the alpha power within the active time-window (0.1s to 1.1s after the probe cue onset) was reduced from −0.91 ± 0.45 dB (AAS BCG corrected data) to −0.71 ± 0.29 dB (non-BCG corrected data) in the Oz electrode, revealing that the BCG artefacts, which are typically below 20Hz, masked the alpha power change related to the visual stimuli. In Figure 11, the cluster-based permutation tests demonstrated significant differences (p < .05) between the AAS BCG corrected and non-BCG corrected data in the occipital electrodes (O1, Oz, O2), confirming that the visual alpha ERD can be observed at the occipital electrodes in the standard AAS BCG corrected data, when compared to the non-BCG corrected data at the sensor level.

**Figure 10.**
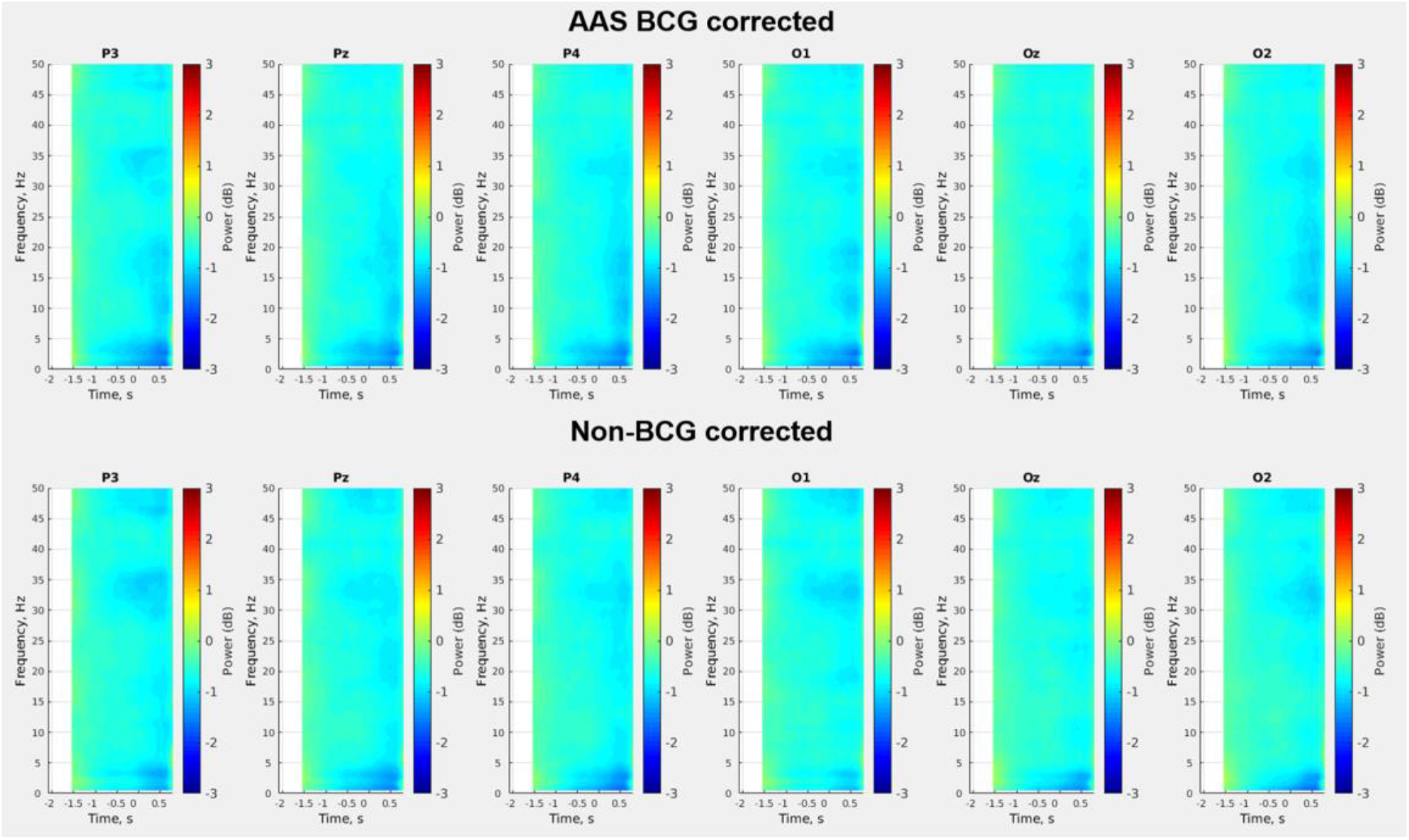
Group mean (N = 20) time-frequency representations (TFRs) of visual alpha (8-13Hz) event-related desynchronization (ERD) in response to the target cues which occurred at 600ms after the probe cue onset (either no cue, double cue, or spatial cue) from 6 electrodes in the occipital and parietal regions for the AAS BCG corrected data (top row) and non-BCG corrected data (bottom row). These TFRs demonstrated −2s to 0.75s relative to the probe cue onset removing a period of the motion artefacts caused by the button press for the task responses occurring at around 1s. The whole trial epoch (−2s to 3s relative to the probe cue onset) can be seen in Supplementary figure (Figure S2).

**Figure 11.**
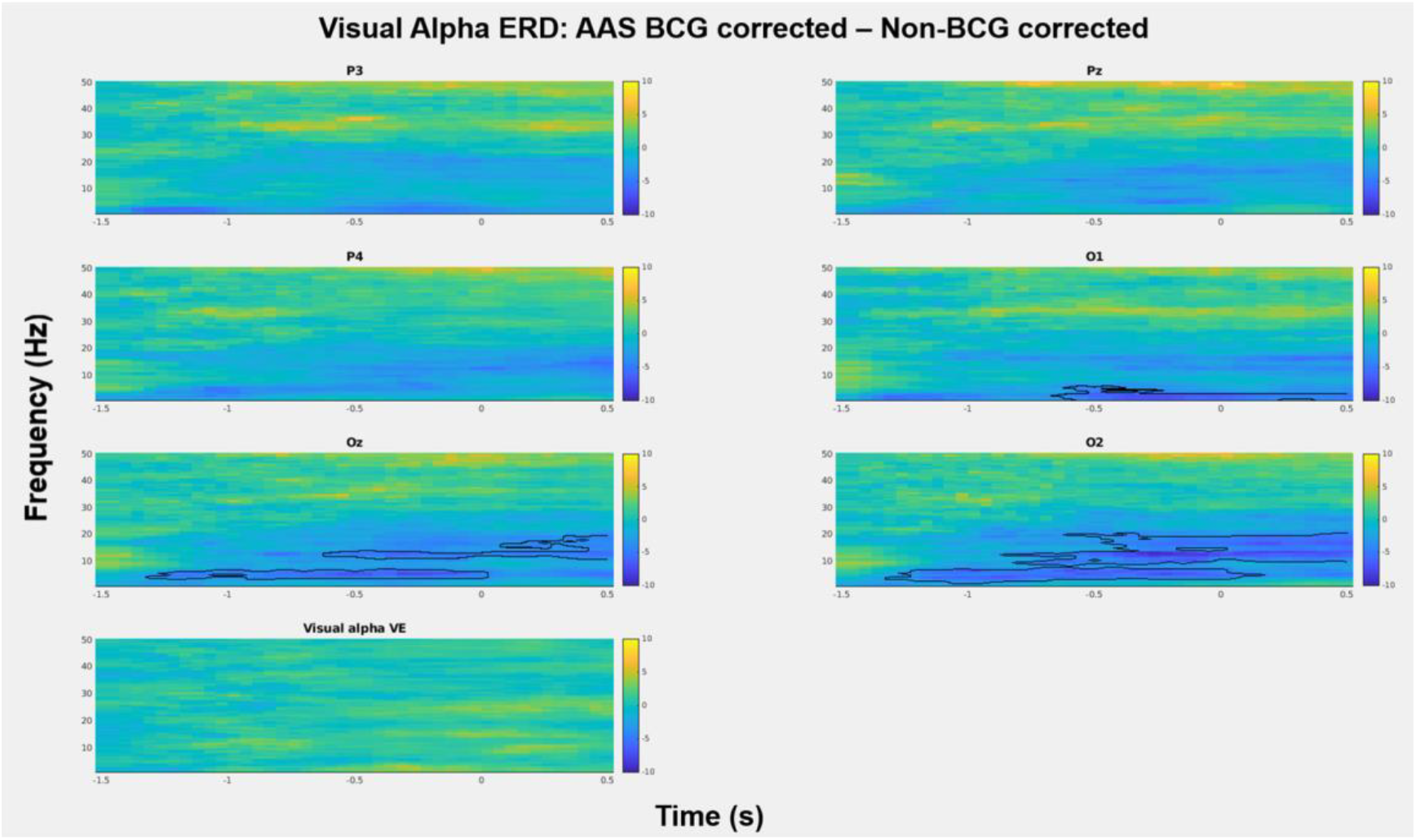
Alpha ERD induced during ANT: The t-maps of differences in the time-frequency representations (TFRs) between the AAS BCG corrected and non-BCG corrected data at 6 electrodes selected from the occipital and parietal regions, and visual alpha VE. Figures demonstrate t-values and black contour representing the significant clusters from cluster-based permutation tests (p < .05), suggesting that AAS BCG corrected data were able to identify a significantly larger ERD decrease in alpha power, when compared to non-BCG corrected data on O1, Oz and O2 electrodes. Yellow denotes a positive and blue denotes a negative t-values.

##### Beamforming source reconstructions

Figure 7 (left column) shows the group average T-statistic map of alpha power changes during the active time-window (0.1s to 1.1s after the probe cue onset) compared to the baseline time-window (−1.5s to −0.5s prior to the probe cue onset) from the BCG corrected and non-BCG corrected data. Alpha power decrease (ERD, negative T values, Figure 7) was observed in the visual cortex in both beamforming+AAS BCG corrected and beamforming BCG corrected datasets. Specifically, the group mean visual alpha VE locations across subjects (±SE) was found at [−25±4, −75±1, −8±3] mm [MNI:x,y,z] for the beamforming AAS BCG corrected data and [2±4, −96±1, 9±3] mm [MNI:x,y,z] for the beamforming BCG corrected data (see Figure 7, crosshair).

Figure 14 (left column) shows the group average TFRs measured in the visual alpha VE for alpha (8-13Hz) power change in response to the target cues which occurred at 600ms after the probe cue onset (either no cue, double cue, or spatial cue) from the AAS BCG corrected data (top row) and non-BCG corrected data (bottom row). Although the broadband increases in power (red vertical stripes occurred after around 1s in response to the probe cue onset) was caused by motion artefacts occurring at the time of the button press for the task responses, alpha power decrease (alpha ERD) was observed within the time window before 1s as compared to baseline time-window (−1.5s to −0.5s prior to the probe cue onset) in both AAS BCG corrected and non-BCG corrected data (see left column in Figure 14). The alpha ERD within the active time-window (0.1s to 1.1s after the probe cue onset) from the BCG corrected data was −1.26 ± 0.48 dB (see top row in Figure 14), whereas the alpha ERD from the non-BCG corrected data was −1.32 ± 0.56 dB (see bottom row in Figure 14). In Figure 15b, the cluster-based permutation tests revealed significant differences (p < .05) between the beamforming BCG corrected data (visual alpha VE) and corresponding non-BCG corrected electrodes (Oz), indicating that the beamforming technique better recovered the meaningful task induced activity (visual alpha ERD) which was not present at the sensor level.

By comparing the standard AAS approach (AAS BCG corrected data) and beamforming+AAS BCG corrected data (Figure 15a), the cluster-based permutation tests revealed significant differences (p < .05) between the beamforming+AAS BCG corrected data (visual alpha VE) and corresponding AAS BCG corrected electrodes (Oz), suggesting the beamforming spatial filtering could improve the SNR of the meaningful task induced activity (visual alpha ERD), when compared to the standard AAS approach in the sensor space. Furthermore, the cluster-based permutation tests (Figure 15c) revealed significant differences (p < .05) between the beamforming BCG corrected data (visual alpha VE) and corresponding AAS BCG corrected electrodes (Oz), demonstrating that the task induced activity of this beamforming BCG denoising data (Figure 14) had significantly greater SNR when compared to that of the conventional AAS approach (Figure 10). By comparing the task induced activity of both beamforming approaches (beamforming+AAS BCG corrected, beamforming BCG corrected), the permutation tests revealed no significant difference between them in the visual alpha VE (Figure 11).

##### Signal to noise ratios (SNRs)

The group mean SNR of the visual alpha VE from the beamforming non-BCG corrected data was 1.03 ± 0.42 (mean ± SD), whereas the group mean SNR of the visual alpha VE from the beamforming+AAS BCG corrected data was 0.99 ± 0.40 (mean ± SD). Both SNRs obtained the source level after beamforming spatial filtering was more than twice as large as the SNRs in the occipital and parietal electrodes ranging between 0.30 and 0.43 in the sensor level of the BCG corrected signals (see Figure 16), which is consistent to the permutation test results.

#### Motor beta event-related desynchronization (ERD)

##### Analysis at the sensor level

Figure 12 shows the group average TFRs measured in the central electrodes (frontal, central, & parietal) for beta (13-30Hz) power change during the motor preparation/planning before the movement onset for the AAS BCG corrected data (top row) and non-BCG corrected data (bottom row). Although the broadband increases in power (red vertical stripes occurred at the time of 0s which was the button press for the task responses) revealing the motion artefacts caused by the button press for the task responses can be observed (see Supplementary Figure: S3), beta power decrease (beta ERD) was observed within the time window before the movement onset as compared to baseline time-window (−3s to −2s prior to the movement onset) in the AAS BCG corrected data (see top row in Figure 12). However, this beta ERD was difficult to identify in the same electrodes from the non-BCG corrected data (see bottom row in Figure 12) as the beta power within the active time-window (−1.25s to −0.25s before the movement onset) was reduced from −0.92 ± 0.23 dB (AAS BCG corrected data) to −0.78 ± 0.25 dB (non-BCG corrected data) in the C4 electrode and from −1.03 ± 0.21 dB (AAS BCG corrected data) to −0.91 ± 0.30 dB (non-BCG corrected data) in the P4 electrode, revealing that the BCG artefacts, which are typically below 20Hz, were masking the beta power change related to the movement preparation/planning. In Figure 13, the cluster-based permutation tests demonstrated that significant differences (p < .05) between the AAS BCG corrected and non-BCG corrected data were observed in the frontal, central, and parietal electrodes (F4, C4, P4) particularly in the right hemisphere which is contralateral to the left-hand button press, confirming that the motor beta ERD was visible at the contralateral frontal, central, and parietal electrodes in the standard AAS BCG corrected data when compared to the non-BCG corrected data at the sensor level.

**Figure 12.**
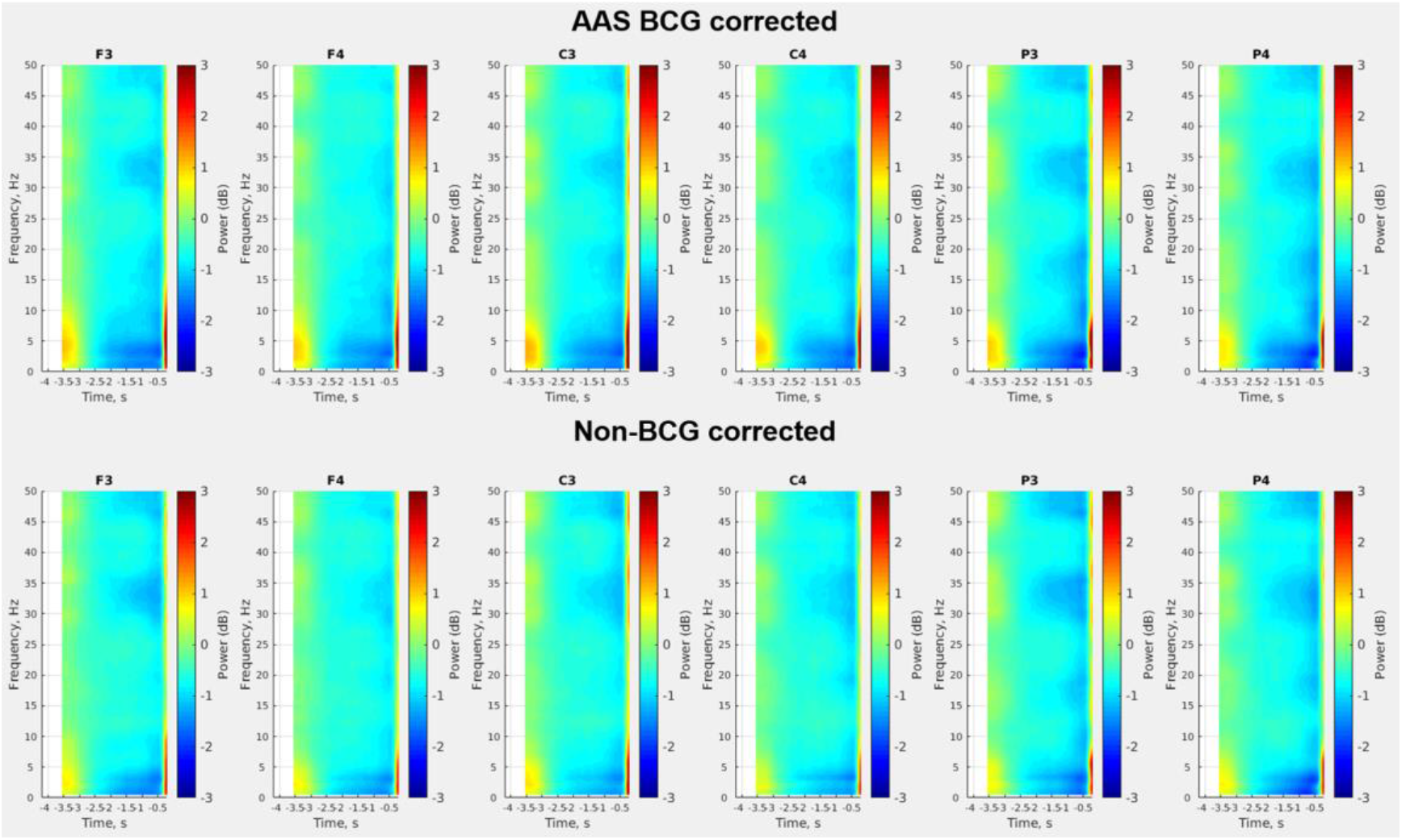
Group mean (N = 20) time-frequency representations (TFRs) of motor beta (13-30Hz) event-related desynchronization (ERD) during motor preparation/planning before the movement onset from 6 electrodes in the frontal, central and parietal regions for the AAS BCG corrected data (top row) and non-BCG corrected data (bottom row). These TFRs demonstrated −4s to −0.25 s relative to the movement onset removing a period of the motion artefacts caused by the button press for the task responses occurring at 0s. The whole trial epoch (−4s to 2s relative to the movement onset) can be seen in Supplementary figure (Figure S3).

**Figure 13.**
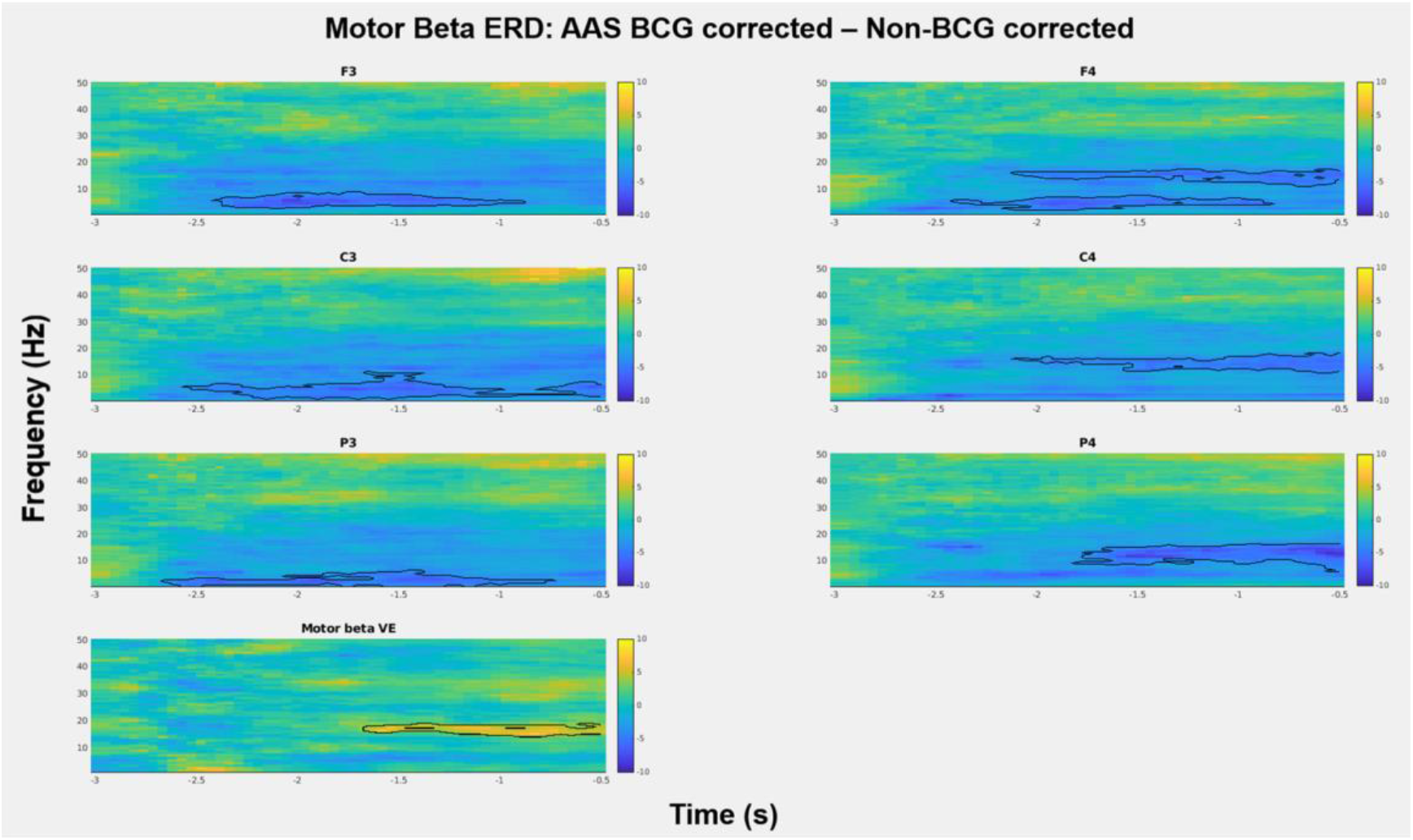
Beta ERD induced during ANT: The t-maps of differences in the time-frequency representations (TFRs) between the AAS BCG corrected and non-BCG corrected data at 6 electrodes selected from the frontal, central and parietal regions, and motor beta VE. Figures demonstrate t-values and black contour representing the significant clusters from cluster-based permutation tests (p < .05), suggesting that AAS BCG corrected data were able to identify a significantly larger ERD decrease in beta power, when compared to non-BCG corrected data in the frontal, central and parietal electrodes. Furthermore, beamforming BCG corrected data exhibited larger beta ERD than beamforming+AAS corrected data. Yellow denotes a positive and blue denotes a negative t-values.

##### Beamforming source reconstructions

Figure 7 (right column) shows the group average T-statistic map of beta power changes during the active time-window (−1.25s to −0.25s before the movement onset) compared to the baseline time-window (−3s to −2s prior to the movement onset) from the beamforming+AAS BCG corrected and beamforming BCG corrected data. Beta power decrease (ERD, negative T values, Figure 7) was observed in the motor cortex in both data set. Specifically, the mean of the individual subject motor beta VE locations (±SE) was found at [45±5, −44±3, 54±2] mm [MNI:x,y,z] for the beamforming+AAS BCG corrected data and [55±7, −35±3, 50±2] mm [MNI:x,y,z] for the beamforming BCG corrected data, both of which were contralateral to the left hand button presses (see Figure 7, crosshair).

Figure 14 (right column) shows the group average TFRs measured in the motor beta VE for beta (13-30Hz) power change during the motor preparation/planning before the movement onset for the AAS BCG corrected data (top row) and non-BCG corrected data (bottom row). Although the broadband increases in power (red vertical stripes occurred at the time of 0s which was the button press for the task responses) revealed the motion artefacts caused by the button press for the task responses, beta power decrease (beta ERD) was observed within the time window before the movement onset as compared to baseline time-window (−3s to −2s prior to the movement onset) in both AAS BCG corrected and non-BCG corrected data (right column in Figure 14). The beta ERD within the active time-window (−1.25s to −0.25s before the movement onset) from the AAS BCG corrected data was −0.95 ± 0.23 dB (see top row in Figure 14), whereas the beta ERD from the non-BCG corrected data was −1.04 ± 0.21 dB (see bottom row in Figure 14). In Figure 15b, the cluster-based permutation tests revealed significant differences (p < .05) between the beamforming BCG corrected data (motor beta VE) and corresponding non-BCG corrected electrodes (C4), indicating that the beamforming technique recovered the meaningful task induced activity (motor beta ERD) which was not present at the sensor level.

**Figure 14.**
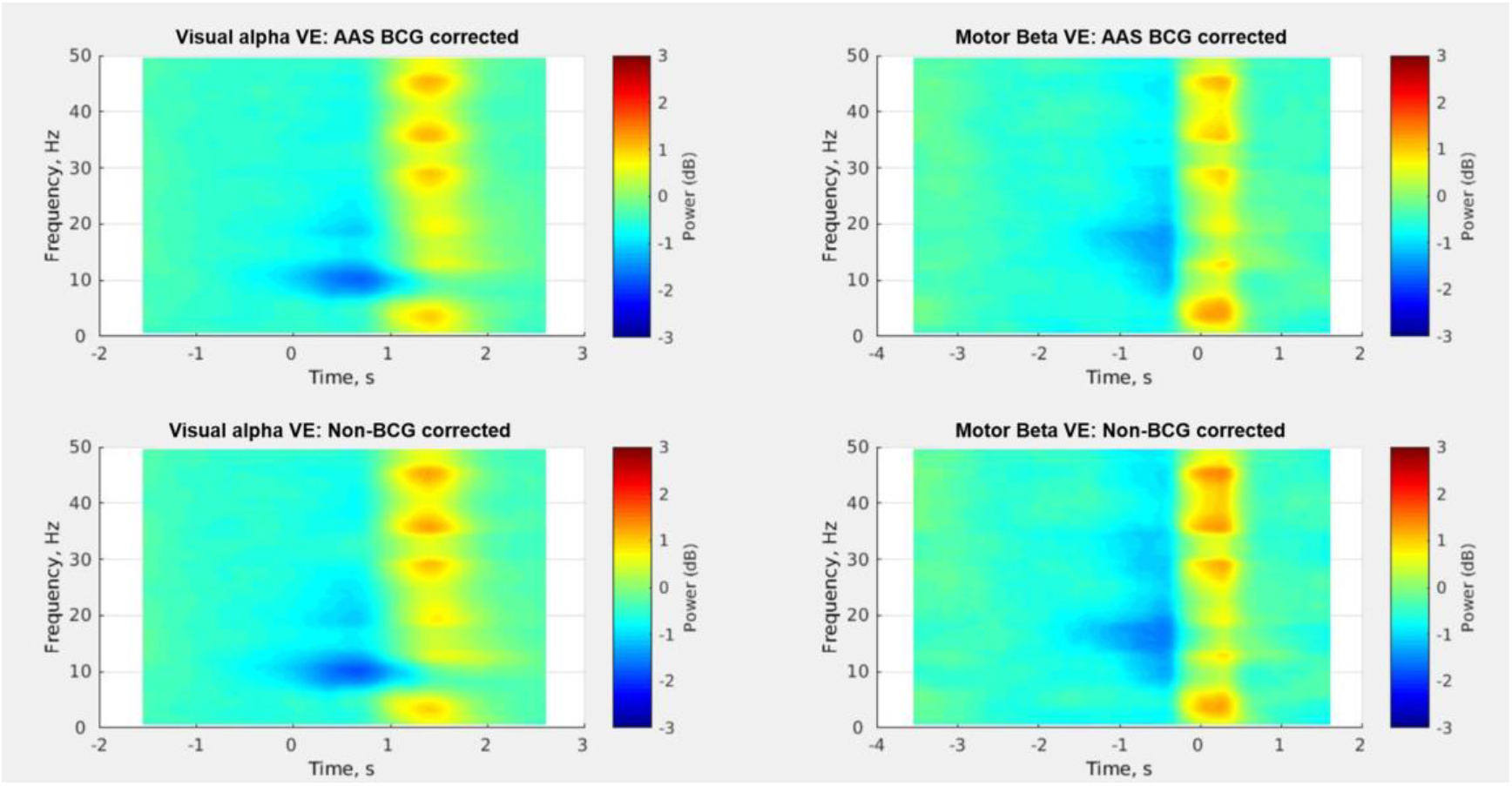
Group mean (N = 20) time-frequency representations (TFRs) of visual alpha (8-13Hz) event-related desynchronization (ERD) in response to the target cues which occurred at 600ms after the probe cue onset (either no cue, double cue, or spatial cue) from the visual alpha VE location for the AAS BCG corrected data (top row) and non-BCG corrected data (bottom row) in the left column, whereas group mean (N = 20) time-frequency representations (TFRs) of motor beta (13-30Hz) event-related desynchronization (ERD) in response to the movement onset from the motor beta VE location for the AAS BCG corrected data (top row) and non-BCG corrected data (bottom row) in the right column. The broadband increases in power (yellow-red vertical stripes) revealed the motion artefacts caused by the button press for the task responses.

**Figure 15.**
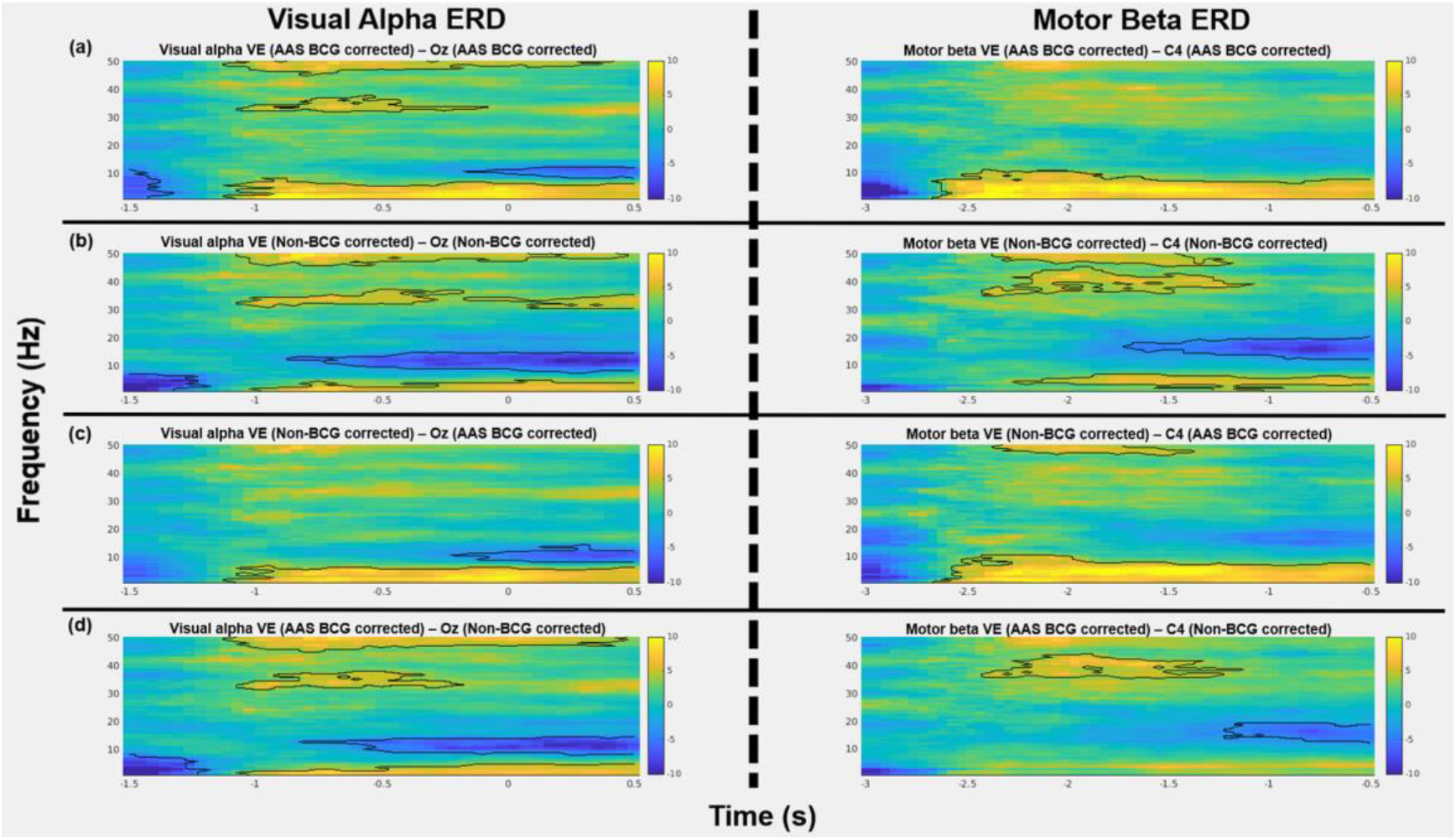
Alpha and Beta ERD induced during ANT: The t-maps of differences between the time-frequency representations (TFRs) for various comparisons, (a) between beamforming+AAS BCG corrected and corresponding AAS BCG corrected EEG electrodes, (b) between beamforming BCG corrected and corresponding non-BCG corrected EEG electrodes, (c) between beamforming BCG corrected and corresponding AAS BCG corrected EEG electrodes, (d) between beamforming+AAS BCG corrected and corresponding non-BCG corrected EEG electrodes, in the visual alpha ERD in the left column and the motor beta ERD in the right column. Figures demonstrate t-values and black contour representing the significant clusters from cluster-based permutation tests (p < .05). Yellow denotes a positive and blue denotes a negative t-values.

By comparing with the standard AAS BCG corrected data, the cluster-based permutation tests revealed significant differences (p < .05) between the beamforming+AAS BCG corrected data (motor beta VE) and corresponding AAS BCG corrected electrodes (C4), as well as significant differences (p < .05) between the beamforming BCG corrected data (motor beta VE) and corresponding AAS BCG corrected electrodes (C4), indicating that both beamforming approaches data (Figure 14) increased SNR significantly when compared to that of the conventional AAS approach (Figure 12). By comparing the task induced activity of both beamforming approaches (beamforming+AAS BCG corrected, beamforming BCG corrected), the permutation test revealed a significant difference (p < .05) between them in the motor beta VE (Figure 13), demonstrating that the beamforming BCG corrected data had significantly larger beta ERD than the beamforming+AAS BCG corrected data.

##### Signal to noise ratios (SNRs)

The group average SNR of the motor beta VE from the non-BCG corrected data was 1.31 ± 0.29 (mean ± SD), whereas the group mean SNR of the motor beta VE from the AAS BCG corrected data was 1.21 ± 0.32 (mean ± SD). Both SNRs in the source level were more than 1.3 times as large as the SNRs in the sensor level for frontal, central and parietal electrodes, ranging between 0.68 and 0.88 in the sensor level of the AAS BCG corrected signals (see Figure 16), which is consistent to the permutation test results.

**Figure 16.**
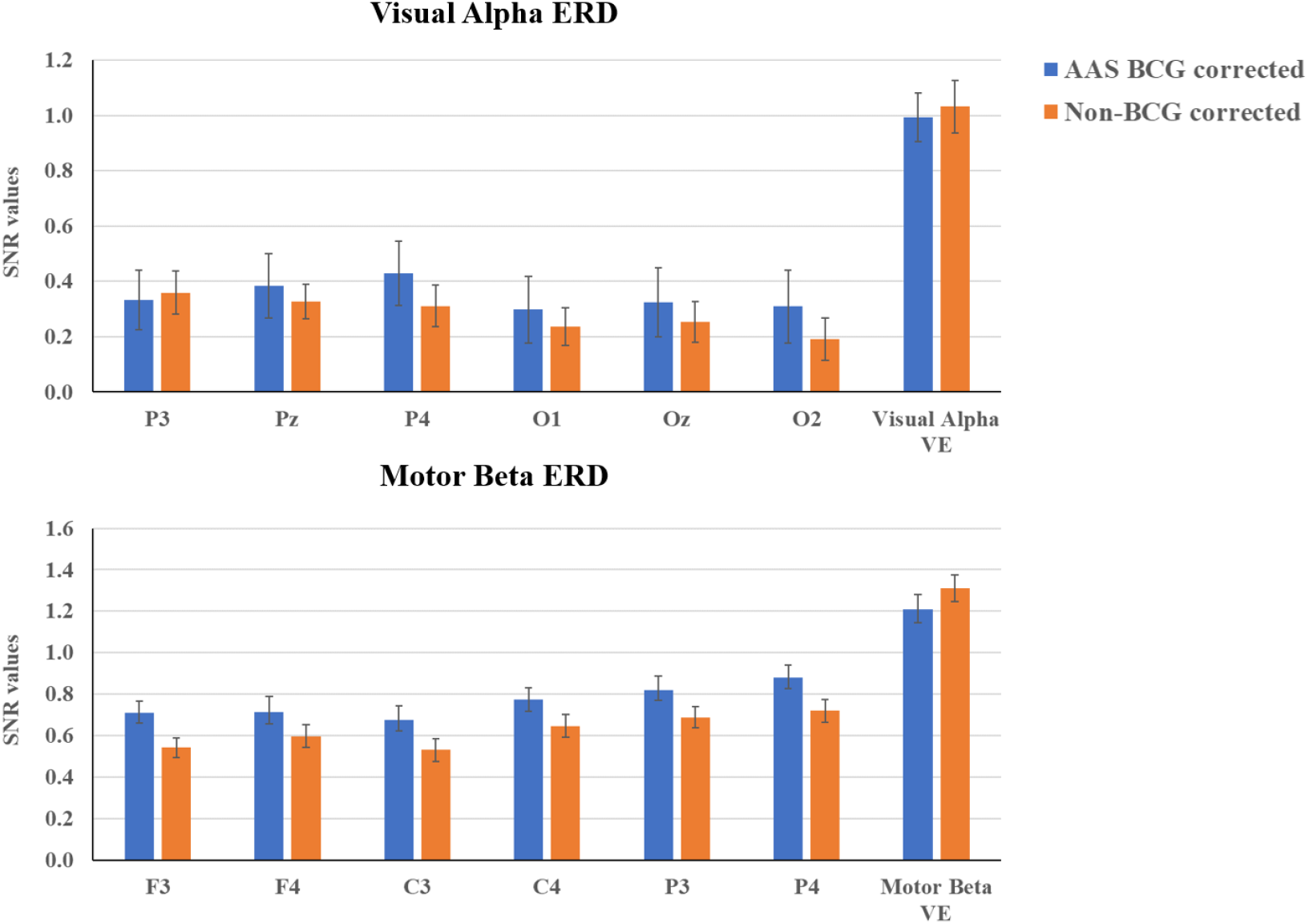
Group mean (N = 20) signal to noise ratios (SNRs) (± SD) of visual alpha (8-13Hz) event-related desynchronization (ERD) for the 6 electrodes of occipital and parietal regions and visual alpha VE location from both AAS BCG corrected and non-BCG corrected data in the top row, and group mean (N = 20) SNRs (± SE) of motor beta (13-30Hz) ERD for the 6 electrodes of frontal, central, and parietal regions and motor beta VE location from both AAS BCG corrected and non-BCG corrected data in the bottom row. SNRs were calculated by dividing mean signal in the frequency of interest within the active time window by the standard deviation of signals in the frequency of interest within the baseline time window, then group mean SNR was calculated averaging averaged individual subject’s SNRs across trials (in a similar approach to Hill et al. (2020)).

## Discussion

In this study, we first investigated the effect of beamforming techniques to attenuate ballistocardiogram (BCG) artefacts in EEG-fMRI, even without detecting cardiac pulse (R-peak) events from simultaneous ECG recordings. In a second step, we also quantified how this technique would preserve expected brain activity induced by the ANT task, while suppressing the BCG artefacts. The beamforming technique is a spatial filtering technique, considered as a dipole scanning approach, proposed for source localization of EEG and MEG data (Gross et al., 2001; Robinson and Vrba, 1999; Sekihara et al., 2001; van Drongelen et al., 1996; van Veen et al., 1997; van Veen and Buckley, 1988). Moreover, several studies have demonstrated that beamformer is highly efficient when attenuating artefactual signals which have different spatial origins from the underlying signal of interest such as eye movements (Cheyne et al., 2006) and orthodontic metal braces (Cheyne et al., 2007) in MEG. Specifically, the beamforming spatial filter rejects sources of signal variance that are not concordant in predetermined source locations in the brain based on the forward model, and attenuates all unwanted source activities outside of the predetermined source location of interest in the data (e.g., eye movements) without having to specify the location or the configuration of these unwanted underlying source signals (Brookes et al., 2007; Huang et al., 2004; van Veen et al., 1997). Since the beamforming spatial filtering appears another promising denoising technique in the context of EEG-fMRI studies (Brookes et al., 2009, 2008a; Mullinger and Bowtell, 2011), we hypothesized that the beamforming technique would attenuate the BCG artefacts to a similar extent as compared to the conventional AAS BCG artefact corrections.

### BCG corrections in EEG-fMRI

We first demonstrated, in the sensor space using time-course and tine-frequency analysis, that the standard AAS method, in general, demonstrated significant suppression of BCG artefacts by comparing the AAS BCG corrected and non-BCG corrected data, while some of the residual BCG artefacts were still observed above 20Hz. We then evaluated the impact of beamforming spatial filtering, when applied to both AAS BCG corrected (beamforming+AAS BCG corrected) and non-BCG corrected (beamforming BCG corrected) data, comparing with sensor level data (AAS BCG corrected data).

The virtual electrode (VE) locations were determined by the ANT task activity, resulting in one VE in the visual cortex (Visual Alpha VE) to monitor alpha power decrease induced by visual cues, and one VE in the motor cortex (Motor Beta VE) to monitor beta power decrease induced by finger tapping preparation/planning. The location of these VEs was estimated specifically for each subject and was found overall concordant at the group level. Based on the two VE locations, we demonstrated that the VE time-courses from both beamforming approaches (beamforming BCG corrected; beamforming AAS+BCG corrected) contained less BCG artefacts when compared to the original uncorrected data in the sensor level. Moreover, both beamforming approaches revealed significantly less BCG artefacts as compared to the standard AAS BCG corrected data. Furthermore, when applying the beamforming to the AAS BCG corrected data, this approach significantly improved the BCG artefact suppressions even after the standard AAS approach, and this approach was significantly better than solely applying the beamforming to the uncorrected data. Overall, our findings demonstrated that the denoising performance of the beamforming BCG corrections was significantly better than the level of the standard AAS approach. Our results support and extend the previous findings of Brookes et al. (2008) which had a limited number of participants (N=2) and did not use any statistical analysis. Our findings revealed that this data-driven beamforming denoising approach even without detecting R-peak events from ECG recording is promising when attenuating the BCG artefacts in EEG-fMRI.

Although the AAS BCG corrections are commonly used and reported in the EEG-fMRI literature, there are several different approaches to correct the BCG artefacts considering both software (Independent Component Analysis (ICA): Debener et al., 2007; Optimal Basis Set (OBS): Niazy et al., 2005) and hardware (Reference layer artefact subtraction: Chowdhury et al., 2014; Carbon-wire loop: Masterton et al., 2007; Optical Motion tracking system: LeVan et al., 2013) based solutions. In the software solutions, the ICA technique has been suggested to remove independent components (ICs) corresponding to the BCG artefacts. In the context of the ICA approaches, several different approaches have been suggested, and the selection of the ICs varies based on 1) thresholding the correlation coefficients (i.e. > 0.2) between each IC and the simultaneously acquired ECG signal (Mantini et al., 2007) or a pulse artefact/BCG template (Srivastava et al., 2005), 2) applying the autocorrelation function of each IC to find a repeating pattern in EEG (Vanderperren et al., 2010), 3) identifying ICs that exhibit spectral peaks at cardiac-related frequencies (Vanderperren et al., 2007), 4) thresholding averaged peak-to-peak amplitudes of ICs reconstructed signals (i.e. 15% and 25%) when epoched around the cardiac peak (Vanderperren et al., 2010), 5) identifying ICs explaining the largest amount of variance of the evoked BCG artefacts (Debener et al., 2008). Although the ICA based approach demonstrated successful BCG suppressions, this method needs more fined parameter tuning. Therefore, in the ICA approaches, the objective and accurate classifications/selections of BCG-related ICs always remain a major concern (Abreu et al., 2016; Vanderperren et al., 2010).

The OBS approach is another software solution, which uses principal component analysis (PCA) to remove the principal components (PCs) representing the BCG artefacts. The OBS relies on the assumption that each BCG artefact occurrence in each channel is independent of any previous occurrence, and Optimal Basis Set (OBS) of principal components is used for the template creation and subsequent BCG artefact subtractions (Niazy et al., 2005). In comparison to ASS, PCA considered in OBS will allow modelling some variability in the BCG artefact, as opposed to considering the mean effect only. In the OBS approach, one parameter needs to be determined, which is the number of principal components used to create the artefact template. Most studies use the recommended value of 3 or 4 for this parameter (Niazy et al., 2005; Vanderperren et al., 2010), although there is no consensus on whether this number can yield satisfactory corrections for all channels of all datasets (Jorge et al., 2019; Marino et al., 2018; Vanderperren et al., 2010). As different from the ICA approach, the OBS method requires the detections of cardiac peak timing similarly to the AAS approach. Although both ICA and OBS approach have successfully removed the BCG artefacts, using some arbitrary classifications/selections in necessary parameters of both approaches remains a major concern, since both approaches have some risk of aggressively suppressing the meaningful signals too much (Abreu et al., 2016; Vanderperren et al., 2010). Moreover, the common application of the ICA and OBS approach have been proposed after the AAS BCG corrections to remove the residual BCG artefacts (Abreu et al., 2018).

Considering these software solutions (ICA & OBS), the AAS approach has still been commonly used and reported in the EEG-fMRI literature. Additionally, this approach has still been recognized as the standard approach (Abreu et al., 2018). However, this AAS approach requires high precisions of the cardiac pulse (R-peak) event detections in the MRI scanner which are used for subtracting averaged BCG artefact templates. Normally R-peak events are detected from simultaneous electrocardiography (ECG) recordings. To make this harder, ECG signals in the MRI scanner are often distorted, which may prove to be even more problematic (Chia et al., 2000; Mullinger et al., 2008b), and significantly time-consuming when manual correction is required. Detecting R-peak events from the ECG is thus difficult, so that this procedure may sometimes become unreliable, although vectorcardiogram (VCG) instead of ECG recordings are more suited and recommended to use for the R-peak detections if available (Mullinger and Bowtell, 2011) because the VCG signals are not distorted as compared to the ECG signals in the MRI. Despite known limitations of the AAS approach, the AAS approach has been used as some ground truth in EEG-fMRI to demonstrate the denoising performance of new techniques. Considering the difficulty and imprecise R peak detections, our proposed data-driven beamforming denoising technique is promising, demonstrating the advantages of neither relying on ECG signals nor needing to determine some parameters as compared to the other correction techniques (ICA, OBS). In this study, solely applying the beamforming spatial filtering technique to the uncorrected data demonstrated superior denoising performance than the standard AAS BCG correction. This data-driven beamforming technique could be well suited for longer EEG-fMRI data acquisition, such as sleep or resting-state data or epilepsy monitoring, where varieties of cardiac responses would be expected to be large and the risk of losing or saturating ECG signals would be high. Under these conditions, the AAS and other software solutions might not be efficient. Based on our findings, this data-driven approach could be applied to suppress the BCG attracts without having to use the detected R-peak events from ECG recordings during the long acquisition.

### Task-based induced neural activity in EEG-fMRI

Having successfully demonstrated the effect of the beamforming technique to suppress the BCG artefacts, it was also crucial to demonstrate that this technique would not suppress the meaningful EEG signals too much, but also preserve them. In this study, we investigated whether this beamforming technique could recover the meaningful task-based induced neural activities indicated by visual alpha (8-13Hz) event-related desynchronization (ERD) during observing visual stimuli and motor beta (13-30Hz) ERD during movement preparation/planning before the movement onset during ANT (Fan et al., 2007, 2005). First, we demonstrated that the standard AAS BCG corrected data indeed exhibited greater alpha ERD during the visual stimuli in the occipital electrodes, and greater beta ERD during the movement preparation in the central electrodes contralateral to the left hand button press as compared to the non-BCG corrected data. Applying the beamforming techniques to both AAS BCG corrected and non-BCG corrected data, we then demonstrated that both beamforming approaches (beamforming+AAS BCG corrected, beamforming BCG corrected) exhibited greater alpha ERD during the visual stimuli in the visual alpha VE, and greater beta ERD during the movement preparation in the motor beta VE as compared to the non-BCG corrected data. Furthermore, as compared to the AAS BCG corrected data, both beamforming approached demonstrated clearer alpha ERD and beta ERD by suppressing any residual artefacts caused by fMRI acquisition. Considering the effect of the data-driven beamforming approaches, the beamforming BCG corrected data revealed significantly greater motor beta ERD as compared to the beamforming+AAS BCG corrected data, whereas there was no significant difference in the visual alpha ERD. Additionally, the locations of visual alpha VEs in the visual cortex were consistent with the previous findings when the visual stimuli were used to trigger movement executions (Brookes et al., 2005; Scheeringa et al., 2011; Wilson et al., 2019), whereas the locations of motor beta VEs were found as expected in the right motor cortex contralateral to the left-hand button presses (Hill et al., 2020; Wilson et al., 2019). Based on these robust visual alpha ERD and motor beta ERD and VE locations, our findings revealed that our data-driven beamforming approaches did not only suppress the BCG artefacts, but also recovered meaningful task-based neural signals induced by the ANT task (Fan et al., 2007; Marshall et al., 2015).

### Signal to noise ratios (SNRs) between sensor and source space

Findings of the visual alpha and motor beta ERDs together demonstrated both beamforming approaches (beamforming+AAS BCG corrected & beamforming BCG corrected data) successfully recovered the expected task-based induced neural activity during visual stimuli or preparation/planning of the movement execution respectively, while minimizing the effects of the BCG artefacts. Furthermore, SNR analysis revealed that the group mean SNRs of the visual alpha VEs from both beamforming approaches in the source level increased the SNRs by more than twice when compared to the SNR measured at the sensor level. The group mean SNRs of the motor beta VEs from both beamforming approaches in the source level increased the SNRs by more than 1.3 times as compared to those in the sensor level, which are consistent to the permutation test results in the TFRs.

Indeed, in a recent MEG study, Hill et al (2020) reported similar beamforming source reconstruction approach improving the SNRs by over 1.5 times for the motor beta frequency activity during finger abduction movements and by over 1.5 times for the visual gamma (55-70Hz) frequency activity during the visual stimuli when compared to those in the sensor space. As expected, our SNR findings were also consistent with the previous findings suggesting that beamforming source reconstruction will improve the SNRs when compared to the sensor level (Brookes et al., 2009; Hill et al., 2020; Sekihara et al., 2004), and this approach will be beneficial especially in the EEG-fMRI (Brookes et al., 2009; Mullinger and Bowtell, 2011), which is indeed the advantage of using beamforming techniques in EEG-fMRI.

Another advantage of applying the beamforming technique in EEG-fMRI is to improve the sensitivity and specificity of fMRI BOLD general linear model (GLM) analysis, although the EEG-fMRI data fusion analysis was beyond the aim of this study. This can be achieved by extracting source activities from a selected VE location, calculating the Hilbert transformed envelope of source neural activities in a specific frequency band, and parametrizing regressors with the Hilbert envelope amplitude fluctuations instead of using consistent fixed binary boxcar (i.e. 0 or 1) for the fMRI GLM analysis (Brookes et al., 2008a; Mullinger et al., 2014). This is based on the findings demonstrating that the amplitudes and variabilities of BOLD responses have been better explained by this parameterized GLM (single-trial correlation analysis) which takes account of source neural variability in EEG responses than by a conventional GLM of consistent amplitude responses (Mullinger et al., 2014). This Hilbert transformed envelope has been used in the different frequency bands (Alpha 8-13Hz: Mullinger et al., 2013, 2014, 2017; Beta 13-30Hz: Pakenham et al., 2020; Wilson et al., 2019; Gamma 55-80Hz: Uji et al., 2018). For example, this parameterized GLM analysis successfully improved the specificity and sensitivity to identify the neurophysiological origin of the negative BOLD response to unilateral median nerve stimulation (Mullinger et al., 2014) and negative BOLD responses between stimulated and unstimulated sensory cortex (Wilson et al., 2019). Furthermore, this GLM analysis revealed that a significant positive correlation between single trial gamma band variability and BOLD responses over the contralateral primary motor cortex during finger abduction movements, indicating that tight coupling of neurovascular coupling between the gamma band activity and BOLD responses (Uji et al., 2018). We assume that such fMRI GLM approach can be used to better localize the neurovascular coupling of sleep specific neural activities (i.e., slow-wave, spindle) in sleep research, after the data-driven beamforming denoising technique suppresses the BCG artefacts which remain problematic for the frequency of sleep specific neural activities.

### Limitations of beamforming techniques

In this paper, we demonstrated that this data-driven beamforming approach did not only significantly attenuate the BCG artefacts without having to use the detected R-peak events from ECG recordings, but also significantly recovered the expected task-based induced brain activity indicated by alpha and beta ERDs when compared to the conventional AAS BCG corrections. However, this beamforming approach also has some limitations. For the construction of the beamforming spatial filters, realistic head volume conductor modelling is required for accurately computing the EEG and MEG lead-fields (Brookes et al., 2008a; Neugebauer et al., 2017). The limitation of the beamforming technique comes from the difficulty in accurately calculating the EEG forward solution. More accurate head models based on realistic head geometry obtained from anatomical MR images and knowledge of the accurate location of the EEG electrodes on the head can provide a more accurate forward solution. Whereas access to such information is not always straightforward, in this paper, we did consider a realistic head model, consisting in a 4-layer (scalp, skull, cerebrospinal fluid, and brain) boundary element method (BEM) head model from individual anatomical images, with digitized EEG electrode locations using the EGI GPS to facilitate individualized accurate co-registration of electrode positions with each subject’s anatomical image.

Another limitation is that better suppressions of MRI related artefacts require to use high-density EEG electrodes (at least 64 EEG channels). Brookes et al. (2009) demonstrated that using high-density EEG electrodes improved the level of artefact attenuations increasing SNRs by a factor of around 1.6 if the number of EEG electrodes was increased from 32 to 64 channels when the data was acquired at 7T MRI. Another major limitation of beamforming technique is that this approach cannot properly reconstruct two spatially separate but temporally correlated sources (Brookes et al., 2007; Huang et al., 2004; Quraan and Cheyne, 2010; van Veen et al., 1997). For example, beamforming cancels each other when spatially far from each other (i.e. auditory steady state response) (Brookes et al., 2007) or merge when they are spatially placed close to each other (Huang et al., 2004), although a modified source model instead of a standard single source model allows for beamforming reconstruction of the spatially separate but temporally correlated sources (Brookes et al., 2007).

Another considered important limitation is how to select an accurate VE location in beamforming source space. Future research first needs to investigate what extent selecting an inaccurate VE location would influence the performance of beamforming approach. In most EEG-fMRI studies, the timing of the EEG events/features is considered more important than its localization, when associated to an fMRI GLM analysis (e.g. detection of epileptic discharges, sleep specific discharges). A commonly used approach for the beamforming technique to select the VE location is to identify a maximum/minimum T score value location in the region of interest for any task-based data. Although this approach to select VEs has been used for resting-state data (Hillebrand et al., 2016, 2012), this way might not be optimal in the sleep research due to the propagation of signals of interest and lack of clear baseline. This issue could be solved statistically using bootstrapping techniques and permutation tests of source images. The validation of these approaches first needs to be fully investigated and addressed to improve beamforming approaches. Furthermore, since the beamforming technique has been initially introduced for the source imaging technique in MEG and EEG studies (Gross et al., 2001; Robinson and Vrba, 1999; Sekihara et al., 2001; van Drongelen et al., 1996; van Veen et al., 1997; van Veen and Buckley, 1988), future study needs to investigate the source localization performance (VE selections) of the beamforming approach comparing with any other source imaging technique in the context of EEG-fMRI.

### Implications

Although our data-driven beamforming approach successfully suppressed the BCG artefacts and recovered the expected task-based induced brain activity in a significantly greater extent than the conventional AAS BCG corrections, it could still be better to combine the AAS approach with the beamforming technique if vectorcardiogram (VCG) recording is available (Chia et al., 2000; Mullinger et al., 2008b; Mullinger and Bowtell, 2011) as compared to solely applying it to uncorrected data. The AAS BCG corrections require high precisions of cardiac pulse (R-peak events) detections in the MRI scanner which are used for subtracting averaged BCG artefact templates. Normally R-peak events are detected from simultaneous electrocardiography (ECG) recordings if the VCG is not available. However, ECG signals in the MRI scanner are distorted and have proven to be problematic (Chia et al., 2000; Mullinger et al., 2008b). Our beamforming BCG artefact correction approach is a data-driven method not requiring identifying noise and signal components, so that this approach appears promising, especially for long data acquisition of sleep or resting-state EEG-fMRI. However, from the current data analysis, this approach is not intended to replace the standard MRI related artefact corrections if the VCG is available. Because our findings also demonstrated the beamforming+AAS BCG corrected data contained significantly less remaining BCG artefacts as compared to the beamforming BCG corrected data, the beamforming and accurate AAS approach could be complementary as the combination of both provides better correction than each of them separately.

## Conclusions

In this paper, we demonstrated that our data-driven beamforming approach did not only attenuate the BCG artefacts without having to use the detected R-peak events from ECG recordings, but also recovered the expected task-based induced brain activity indicated by alpha and beta ERDs, which the BCG artefacts typically obscure, in a significantly greater extent than the conventional AAS BCG corrections. Our findings bring new insight into an active area of research in EEG-fMRI related to the extraction of meaningful brain signals and suppression of MRI-related artefacts. This approach would be very promising and beneficial especially for the sleep EEG-fMRI data without detecting R-peak events from ECG recording. Future research will compare this beamforming BCG artefact correction approach with MRI related artefact corrections using VCG recordings, and further examine whether this beamforming approach would be better suited to recover meaningful EEG signals related to spontaneous events such as brain rhythms during sleep and epileptic discharges.

## Acknowledgements

This research was supported by the Natural Sciences and Engineering Research Council of Canada (TDV) and the Canada Foundation for Innovation (TDV). The MRI compatible high-density EEG device (Magstim-EGI) and data acquisition were made possible through an internal grant from PERFORM centre and the Faculty of Arts and Science of Concordia University (CG). TDV is also supported by the Canadian Institutes of Health Research (MOP 142191, PJT 153115, PJT 156125 and PJT 166167), the Fonds de Recherche du Québec – Santé and Concordia University. CG is supported Natural Sciences and Engineering Research Council of Canada Discovery grants as well as the Canadian Institutes of Health Research (PJT-159948 and MOP-133619) and the Fonds de Recherche du Québec, Nature et Technology (research team grant). MU is supported by a Horizon Postdoctoral Fellowship from Concordia University.

## The authorship contribution statement

**Makoto Uji**: Conceptualization, Methodology, Formal data analysis, Investigation, Data curation, Visualization, Writing – original draft, Visualization, Project administration. **Nathan Cross**: Data acquisition, Writing – review & editing. **Florence B. Pomares**: Data acquisition, Writing – review & editing. **Aurore A. Perrault**: Data acquisition, Writing – review & editing. **Aude Jegou**: Data acquisition, Writing – review & editing. **Alex Nguyen**: Data acquisition, Writing – review & editing. **Umit Aydin**: Data acquisition, Writing – review & editing. **Jean-Marc Lina**: Methodology, Validation, Writing – review & editing. **Christophe Grova**: Conceptualization, Resources, Writing – review & editing, Supervision, Project administration, Funding acquisition. **Thien Thanh Dang-Vu**: Conceptualization, Resources, Writing – review & editing, Supervision, Project administration, Funding acquisition.

---

**Figure S1.**
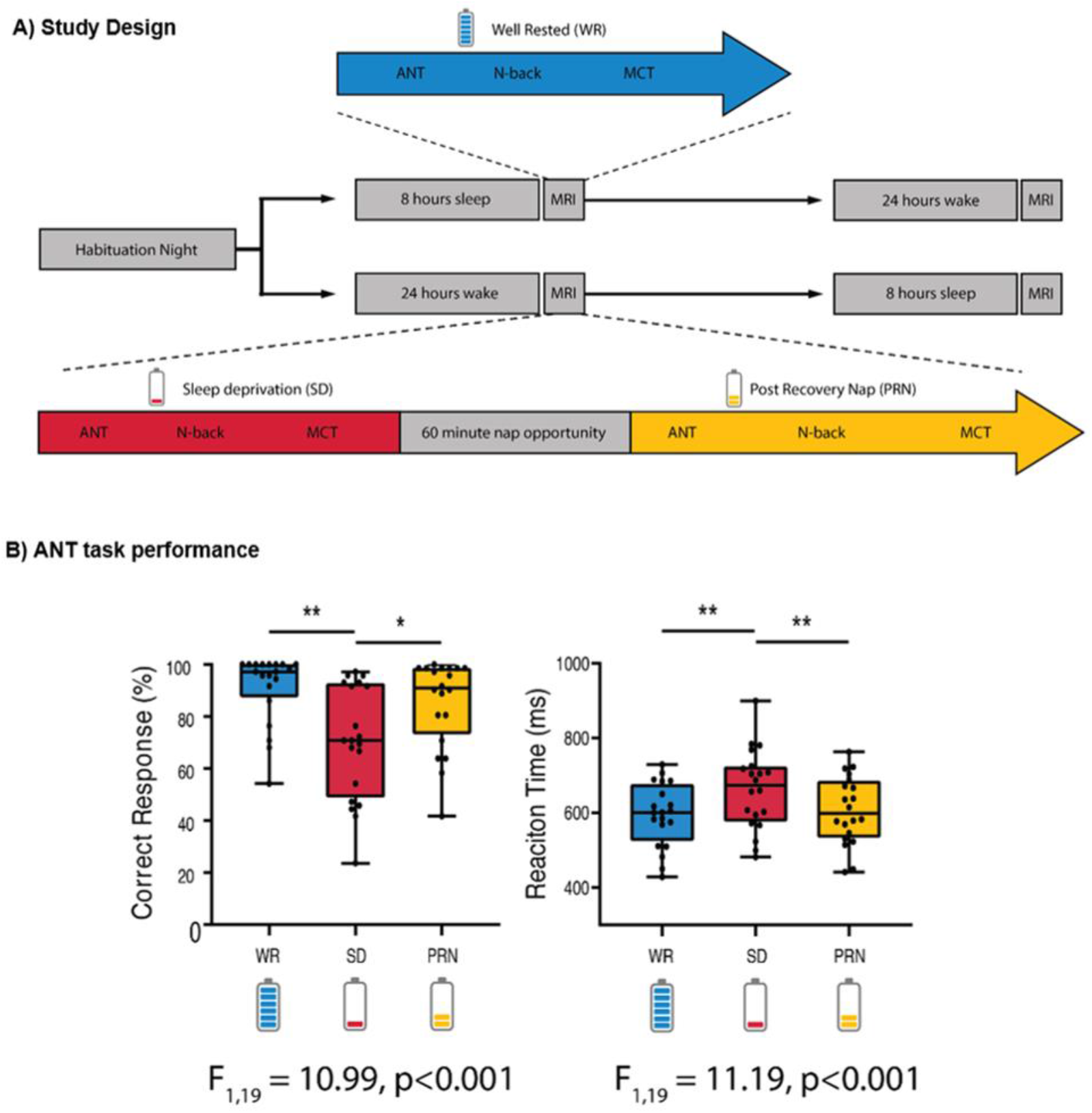
Study design and behavioural results of the attentional network task (ANT). A) Participants made 3 visits to the lab: a habituation night, followed by a counterweighted design of either another full-night opportunity to sleep (blue), or a night of total sleep deprivation. In the morning following each night, participants completed cognitive tasks inside the MRI scanner. In the sleep deprived state (red), participants also had a recovery nap opportunity (yellow) and then completed the tasks inside the MRI scanner again. B) The percentage of correct responses and mean reaction time on the ANT was significantly impaired following sleep deprivation, and improved following a recovery nap.

**Figure S2.**
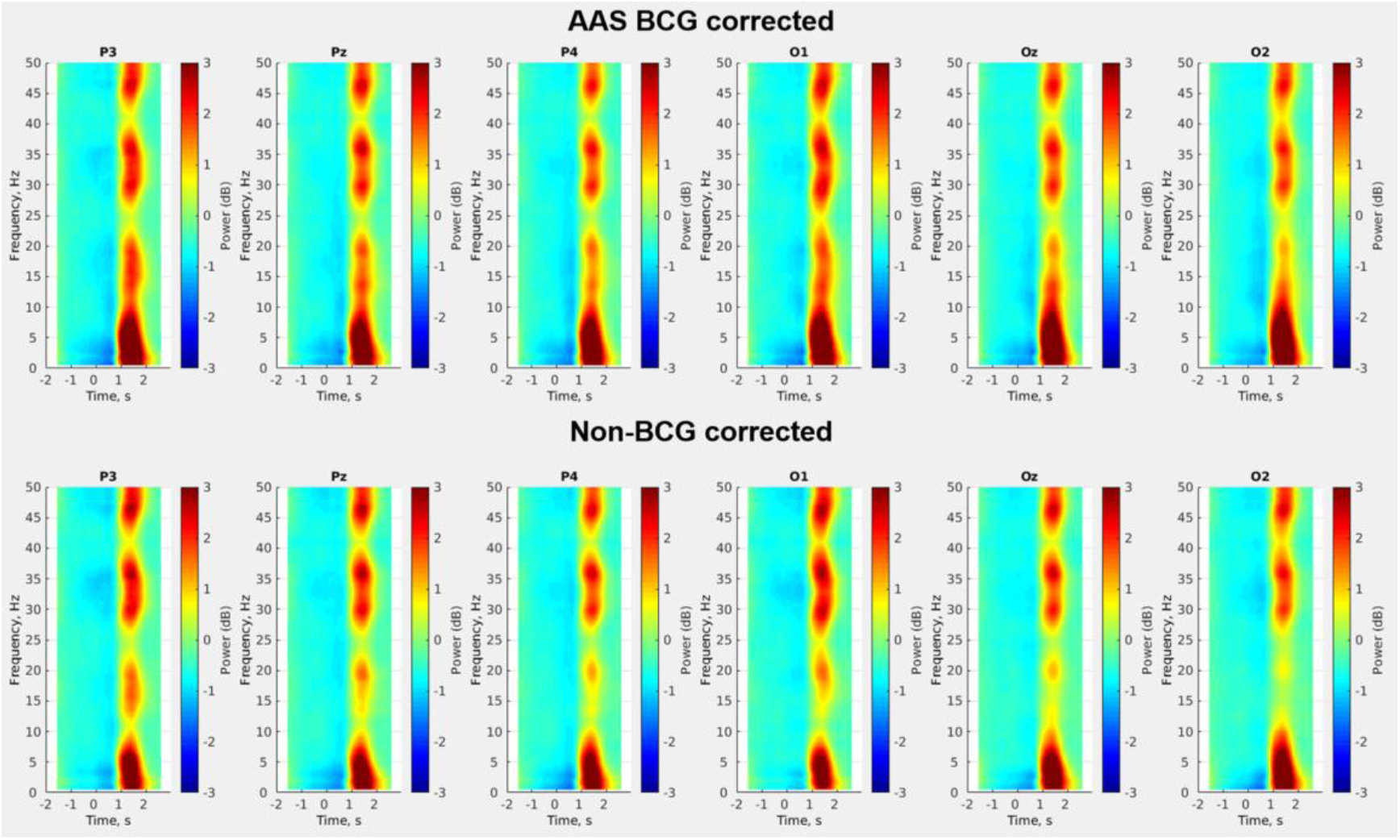
Group mean (N = 20) time-frequency representations (TFRs) of visual alpha (8-13Hz) event-related desynchronization (ERD) in response to the target cues which occurred at 600ms after the probe cue onset (either no cue, double cue, or spatial cue) from 6 electrodes in the occipital and parietal regions for the AAS BCG corrected data (top row) and non-BCG corrected data (bottom row). These TFRs demonstrated the whole trial epoch period (−2s to 3s relative to the probe cue onset). The broadband increases in power (red vertical stripes occurred after around 1s in response to the probe cue onset) revealed the motion artefacts caused by the button press for the task responses.

**Figure S3.**
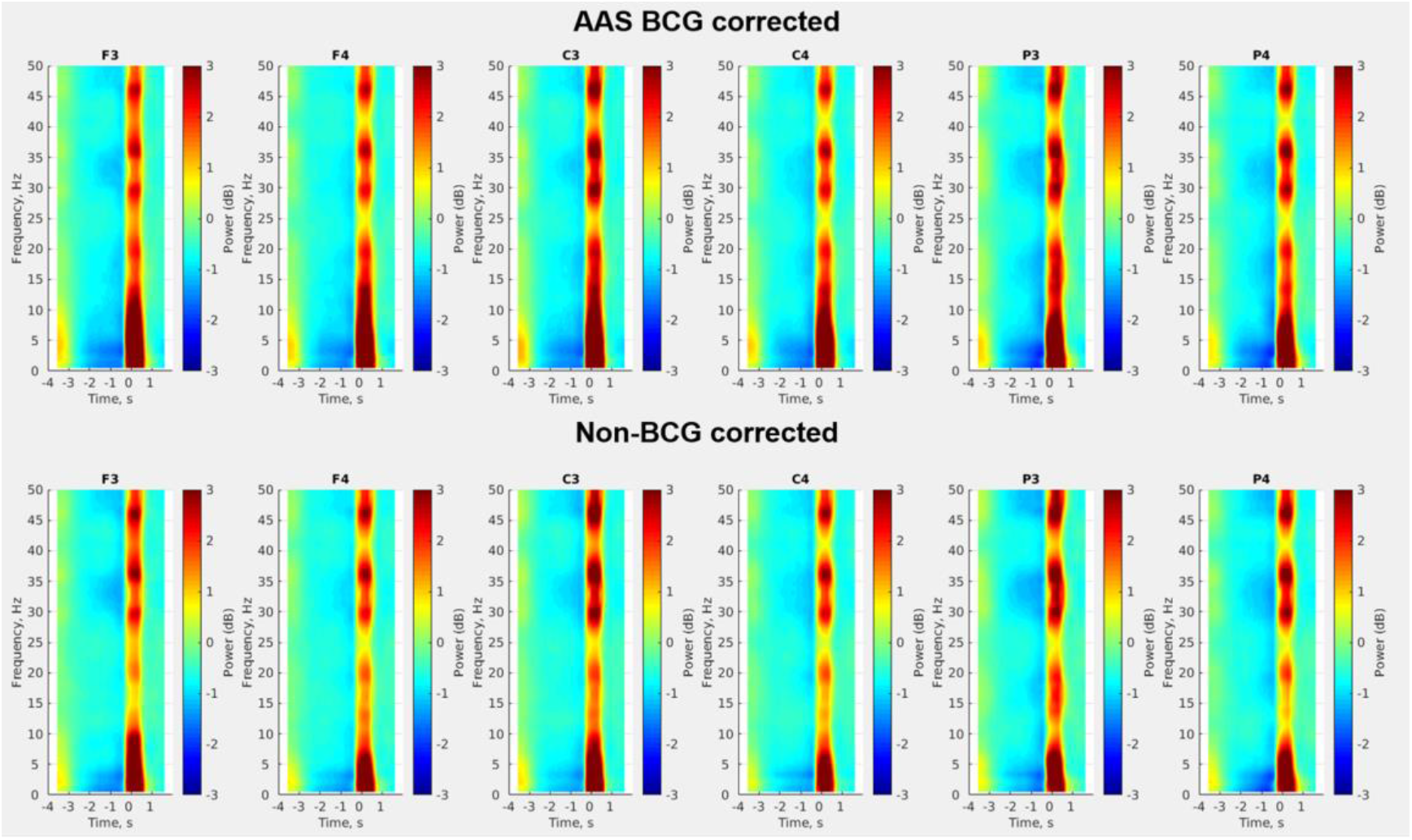
Group mean (N = 20) time-frequency representations (TFRs) of motor beta (13-30Hz) event-related desynchronization (ERD) during motor preparation/planning before the movement onset from 6 electrodes in the frontal, central and parietal regions for the AAS BCG corrected data (top row) and non-BCG corrected data (bottom row). These TFRs demonstrated the whole trial epoch period (−4s to 2s relative to the movement onset). The broadband increases in power (red vertical stripes occurred at 0s in response to the movement onset) revealed the motion artefacts caused by the button press for the task responses.

## Notes

### Competing Interest Statement

The authors have declared no competing interest.

## References

Abreu, R., Leal, A., Figueiredo, P., 2018. EEG-Informed fMRI: A Review of Data Analysis Methods. Front. Hum. Neurosci. 12. https://doi.org/10.3389/fnhum.2018.00029

Abreu, R., Leite, M., Jorge, J., Grouiller, F., van der Zwaag, W., Leal, A., Figueiredo, P., 2016. Ballistocardiogram artifact correction taking into account physiological signal preservation in simultaneous EEG-fMRI. Neuroimage 135, 45–63. https://doi.org/10.1016/j.neuroimage.2016.03.034

Allen, P.J., Josephs, O., Turner, R., 2000. A method for removing imaging artifact from continuous EEG recorded during functional MRI. Neuroimage 12, 230–239. https://doi.org/10.1006/nimg.2000.0599

Allen, P.J., Polizzi, G., Krakow, K., Fish, D.R., Lemieux, L., 1998. Identification of EEG Events in the MR Scanner: The Problem of Pulse Artifact and a Method for Its Subtraction. Neuroimage 8, 229–239. https://doi.org/10.1006/nimg.1998.0361

Bagshaw, A.P., Aghakhani, Y., Bénar, C.G., Kobayashi, E., Hawco, C., Dubeau, F., Pike, G.B., Gotman, J., 2004. EEG-fMRI of focal epileptic spikes: Analysis with multiple haemodynamic functions and comparison with gadolinium-enhanced MR angiograms. Hum. Brain Mapp. 22, 179–192. https://doi.org/10.1002/hbm.20024

Becker, R., Reinacher, M., Freyer, F., Villringer, A., Ritter, P., 2011. How Ongoing Neuronal Oscillations Account for Evoked fMRI Variability. J. Neurosci. 31, 11016–11027. https://doi.org/10.1523/jneurosci.0210-11.2011

Bénar, C.G., Schön, D., Grimault, S., Nazarian, B., Burle, B., Roth, M., Badier, J.M., Marquis, P., Liegeois-Chauvel, C., Anton, J.L., 2007. Single-trial analysis of oddball event-related potentials in simultaneous EEG-fMRI. Hum. Brain Mapp. 28, 602–613. https://doi.org/10.1002/hbm.20289

Bonmassar, G., Purdon, P.L., Jääskeläinen, I.P., Chiappa, K., Solo, V., Brown, E.N., Belliveau, J.W., 2002. Motion and ballistocardiogram artifact removal for interleaved recording of EEG and EPs during MRI. Neuroimage 16. https://doi.org/10.1006/nimg.2002.1125

Brookes, M.J., Gibson, A.M., Hall, S.D., Furlong, P.L., Barnes, G.R., Hillebrand, A., Singh, K.D., Holliday, I.E., Francis, S.T., Morris, P.G., 2005. GLM-beamformer method demonstrates stationary field, alpha ERD and gamma ERS co-localisation with fMRI BOLD response in visual cortex. Neuroimage 26, 302–308. https://doi.org/10.1016/j.neuroimage.2005.01.050

Brookes, M.J., Mullinger, K.J., Stevenson, C.M., Morris, P.G., Bowtell, R., 2008a. Simultaneous EEG source localisation and artifact rejection during concurrent fMRI by means of spatial filtering. Neuroimage 40, 1090–1104. https://doi.org/10.1016/j.neuroimage.2007.12.030

Brookes, M.J., Stevenson, C.M., Barnes, G.R., Hillebrand, A., Simpson, M.I.G., Francis, S.T., Morris, P.G., 2007. Beamformer reconstruction of correlated sources using a modified source model. Neuroimage 34, 1454–1465. https://doi.org/10.1016/j.neuroimage.2006.11.012

Brookes, M.J., Vrba, J., Mullinger, K.J., Geirsdóttir, G.B., Yan, W.X., Stevenson, C.M., Bowtell, R., Morris, P.G., 2009. Source localisation in concurrent EEG/fMRI: Applications at 7T. Neuroimage 45, 440–452. https://doi.org/10.1016/j.neuroimage.2008.10.047

Brookes, M.J., Vrba, J., Robinson, S.E., Stevenson, C.M., Peters, A.M., Barnes, G.R., Hillebrand, A., Morris, P.G., 2008b. Optimising experimental design for MEG beamformer imaging. Neuroimage 39, 1788–1802. https://doi.org/10.1016/j.neuroimage.2007.09.050

Buzsáki, G., Anastassiou, C.A., Koch, C., 2012. The origin of extracellular fields and currents — EEG, ECoG, LFP and spikes. Nat. Rev. Neurosci. 13, 407–420. https://doi.org/10.1038/nrn3241

Cheyne, D., Bakhtazad, L., Gaetz, W., 2006. Spatiotemporal mapping of cortical activity accompanying voluntary movements using an event-related beamforming approach. Hum. Brain Mapp. 27, 213–229. https://doi.org/10.1002/hbm.20178

Cheyne, D., Bostan, A.C., Gaetz, W., Pang, E.W., 2007. Event-related beamforming: A robust method for presurgical functional mapping using MEG. Clin. Neurophysiol. 118, 1691–1704. https://doi.org/10.1016/j.clinph.2007.05.064

Cheyne, D., Ferrari, P., 2013. MEG studies of motor cortex gamma oscillations: evidence for a gamma “fingerprint” in the brain? Front. Hum. Neurosci. 7. https://doi.org/10.3389/fnhum.2013.00575

Chia, J.M., Fischer, S.E., Wickline, S.A., Lorenz, C.H., 2000. Performance of QRS detection for cardiac magnetic resonance imaging with a novel vectorcardiographic triggering method. J. Magn. Reson. Imaging 12, 678–688. https://doi.org/10.1002/1522-2586(200011)12:5<678::AID-JMRI4>3.0.CO;2-5

Chowdhury, M.E.H., Mullinger, K.J., Glover, P., Bowtell, R., 2014. Reference layer artefact subtraction (RLAS): A novel method of minimizing EEG artefacts during simultaneous fMRI. Neuroimage 84, 307–319. https://doi.org/10.1016/j.neuroimage.2013.08.039

Cross, N., Paquola, C., Pomares, F.B., Perrault, A.A., Jegou, A., Nguyen, A., Aydin, U., Bernhardt, B.C., Grova, C., Dang-Vu, T.T., 2021. Cortical gradients of functional connectivity are robust to state-dependent changes following sleep deprivation. Neuroimage 226, 117547. https://doi.org/10.1016/j.neuroimage.2020.117547

Dang-Vu, T.T., Bonjean, M., Schabus, M., Boly, M., Darsaud, A., Desseilles, M., Degueldre, C., Balteau, E., Phillips, C., Luxen, A., Sejnowski, T.J., Maquet, P., 2011. Interplay between spontaneous and induced brain activity during human non-rapid eye movement sleep. Proc. Natl. Acad. Sci. U. S. A. 108, 15438–15443. https://doi.org/10.1073/pnas.1112503108

Dang-Vu, T.T., Schabus, M., Desseilles, M., Albouy, G., Boly, M., Darsaud, A., Gais, S., Rauchs, G., Sterpenich, V., Vandewalle, G., Carrier, J., Moonen, G., Balteau, E., Degueldre, C., Luxen, A., Phillips, C., Maquet, P., 2008. Spontaneous neural activity during human slow wave sleep. Proc. Natl. Acad. Sci. U. S. A. 105. https://doi.org/10.1073/pnas.0801819105

Darvas, F., Scherer, R., Ojemann, J.G., Rao, R.P., Miller, K.J., Sorensen, L.B., 2010. High gamma mapping using EEG. Neuroimage 49, 930–938. https://doi.org/10.1016/j.neuroimage.2009.08.041

Debener, S., Mullinger, K.J., Niazy, R.K., Bowtell, R.W., 2008. Properties of the ballistocardiogram artefact as revealed by EEG recordings at 1.5, 3 and 7 T static magnetic field strength. Int J Psychophysiol 67, 189–199. https://doi.org/10.1016/j.ijpsycho.2007.05.015

Debener, S., Strobel, A., Sorger, B., Peters, J., Kranczioch, C., Engel, A.K., Goebel, R., 2007. Improved quality of auditory event-related potentials recorded simultaneously with 3-T fMRI: removal of the ballistocardiogram artefact. Neuroimage 34, 587–597. https://doi.org/10.1016/j.neuroimage.2006.09.031

Debener, S., Ullsperger, M., Siegel, M., Engel, A.K., 2006. Single-trial EEG-fMRI reveals the dynamics of cognitive function. Trends Cogn. Sci. 10, 558–563. https://doi.org/10.1016/j.tics.2006.09.010

Debener, S., Ullsperger, M., Siegel, M., Fiehler, K., von Cramon, D.Y., Engel, A.K., 2005. Trial-by-trial coupling of concurrent electroencephalogram and functional magnetic resonance imaging identifies the dynamics of performance monitoring. J Neurosci 25, 11730–11737. https://doi.org/10.1523/JNEUROSCI.3286-05.2005

Delorme, A., Makeig, S., 2004. EEGLAB: an open source toolbox for analysis of single-trial EEG dynamics including independent component analysis. J. Neurosci. Methods 134, 9–21. https://doi.org/10.1016/j.jneumeth.2003.10.009

Delorme, A., Sejnowski, T., Makeig, S., 2007. Enhanced detection of artifacts in EEG data using higher-order statistics and independent component analysis. Neuroimage 34, 1443–1449. https://doi.org/10.1016/j.neuroimage.2006.11.004

Eichele, T., Calhoun, V.D., Moosmann, M., Specht, K., Jongsma, M.L.A., Quiroga, R.Q., Nordby, H., Hugdahl, K., 2008. Unmixing concurrent EEG-fMRI with parallel independent component analysis. Int. J. Psychophysiol. 67, 222–234. https://doi.org/10.1016/j.ijpsycho.2007.04.010

Eichele, T., Specht, K., Moosmann, M., Jongsma, M.L.A., Quiroga, R.Q., Nordby, H., Hugdahl, K., 2005. Assessing the spatiotemporal evolution of neuronal activation with single-trial event-related potentials and functional MRI. Proc Natl Acad Sci U S A 102, 17798–17803. https://doi.org/10.1073/pnas.0505508102

Fan, J., Byrne, J., Worden, M.S., Guise, K.G., McCandliss, B.D., Fossella, J., Posner, M.I., 2007. The Relation of Brain Oscillations to Attentional Networks. J. Neurosci. 27, 6197–6206. https://doi.org/10.1523/JNEUROSCI.1833-07.2007

Fan, J., McCandliss, B., Fossella, J., Flombaum, J., Posner, M., 2005. The activation of attentional networks. Neuroimage 26, 471–479. https://doi.org/10.1016/j.neuroimage.2005.02.004

Fan, J., McCandliss, B.D., Sommer, T., Raz, A., Posner, M.I., 2002. Testing the Efficiency and Independence of Attentional Networks. J. Cogn. Neurosci. 14, 340–347. https://doi.org/10.1162/089892902317361886

Fuchs, M., Kastner, J., Wagner, M., Hawes, S., Ebersole, J.S., 2002. A standardized boundary element method volume conductor model. Clin. Neurophysiol. 113, 702–712. https://doi.org/10.1016/S1388-2457(02)00030-5

Fultz, N.E., Bonmassar, G., Setsompop, K., Stickgold, R.A., Rosen, B.R., Polimeni, J.R., Lewis, L.D., 2019. Coupled electrophysiological, hemodynamic, and cerebrospinal fluid oscillations in human sleep. Science (80-.). 366. https://doi.org/10.1126/science.aax5440

Goldman, R.I., Stern, J.M., Engel Jerome, J., Cohen, M.S., 2002. Simultaneous EEG and fMRI of the alpha rhythm. Neuroreport 13, 2487–2492. https://doi.org/10.1097/01.wnr.0000047685.08940.d0

Goldman, R.I., Wei, C.Y., Philiastides, M.G., Gerson, A.D., Friedman, D., Brown, T.R., Sajda, P., 2009. Single-trial discrimination for integrating simultaneous EEG and fMRI: identifying cortical areas contributing to trial-to-trial variability in the auditory oddball task. Neuroimage 47, 136–147. https://doi.org/10.1016/j.neuroimage.2009.03.062

Gotman, J., Kobayashi, E., Bagshaw, A.P., Bénar, C.-G., Dubeau, F., 2006. Combining EEG and fMRI: A multimodal tool for epilepsy research. J. Magn. Reson. Imaging 23, 906–920. https://doi.org/10.1002/jmri.20577

Gotman, J., Pittau, F., 2011. Combining EEG and fMRI in the study of epileptic discharges. Epilepsia 52. https://doi.org/10.1111/j.1528-1167.2011.03151.x

Gramfort, A., Papadopoulo, T., Olivi, E., Clerc, M., 2010. OpenMEEG: opensource software for quasistatic bioelectromagnetics. Biomed. Eng. Online 9, 45. https://doi.org/10.1186/1475-925X-9-45

Gross, J., Kujala, J., Hamalainen, M., Timmermann, L., Schnitzler, A., Salmelin, R., 2001. Dynamic imaging of coherent sources: Studying neural interactions in the human brain. Proc. Natl. Acad. Sci. 98, 694–699. https://doi.org/10.1073/pnas.98.2.694

Grova, C., Daunizeau, J., Kobayashi, E., Bagshaw, A.P., Lina, J.M., Dubeau, F., Gotman, J., 2008. Concordance between distributed EEG source localization and simultaneous EEG-fMRI studies of epileptic spikes. Neuroimage 39. https://doi.org/10.1016/j.neuroimage.2007.08.020

Hale, J.R., White, T.P., Mayhew, S.D., Wilson, R.S., Rollings, D.T., Khalsa, S., Arvanitis, T.N., Bagshaw, A.P., 2016. Altered thalamocortical and intra-thalamic functional connectivity during light sleep compared with wake. Neuroimage 125. https://doi.org/10.1016/j.neuroimage.2015.10.041

Hall, S.D., Stanford, I.M., Yamawaki, N., McAllister, C.J., Ronnqvist, K.C., Woodhall, G.L., Furlong, P.L., 2011. The role of GABAergic modulation in motor function related neuronal network activity. Neuroimage 56, 1506–1510.

Heers, M., Hedrich, T., An, D., Dubeau, F., Gotman, J., Grova, C., Kobayashi, E., 2014. Spatial correlation of hemodynamic changes related to interictal epileptic discharges with electric and magnetic source imaging. Hum. Brain Mapp. 35. https://doi.org/10.1002/hbm.22482

Hill, R.M., Boto, E., Rea, M., Holmes, N., Leggett, J., Coles, L.A., Papastavrou, M., Everton, S.K., Hunt, B.A.E., Sims, D., Osborne, J., Shah, V., Bowtell, R., Brookes, M.J., 2020. Multi-channel whole-head OPM-MEG: Helmet design and a comparison with a conventional system. Neuroimage 219, 116995. https://doi.org/10.1016/j.neuroimage.2020.116995

Hillebrand, A., Barnes, G.R., 2005. Beamformer Analysis of MEG Data. Int. Rev. Neurobiol. https://doi.org/10.1016/S0074-7742(05)68006-3

Hillebrand, A., Barnes, G.R., Bosboom, J.L., Berendse, H.W., Stam, C.J., 2012. Frequency-dependent functional connectivity within resting-state networks: An atlas-based MEG beamformer solution. Neuroimage 59, 3909–3921. https://doi.org/10.1016/j.neuroimage.2011.11.005

Hillebrand, A., Tewarie, P., van Dellen, E., Yu, M., Carbo, E.W.S., Douw, L., Gouw, A.A., van Straaten, E.C.W., Stam, C.J., 2016. Direction of information flow in large-scale resting-state networks is frequency-dependent. Proc. Natl. Acad. Sci. 113, 3867–3872. https://doi.org/10.1073/pnas.1515657113

Horovitz, S.G., Fukunaga, M., De Zwart, J.A., Van Gelderen, P., Fulton, S.C., Balkin, T.J., Duyn, J.H., 2008. Low frequency BOLD fluctuations during resting wakefulness and light sleep: A simultaneous EEG-fMRI study. Hum. Brain Mapp. 29, 671–682. https://doi.org/10.1002/hbm.20428

Huang, M.-X., Shih, J.J., Lee, R.R., Harrington, D.L., Thoma, R.J., Weisend, M.P., Hanlon, F., Paulson, K.M., Li, T., Martin, K., Miller, G.A., Canive, J.M., 2004. Commonalities and Differences Among Vectorized Beamformers in Electromagnetic Source Imaging. Brain Topogr. 16, 139–158. https://doi.org/10.1023/B:BRAT.0000019183.92439.51

Huster, R.J., Debener, S., Eichele, T., Herrmann, C.S., 2012. Methods for simultaneous EEG-fMRI: an introductory review. J Neurosci 32, 6053–6060. https://doi.org/10.1523/JNEUROSCI.0447-12.2012

Ives, J.R., Warach, S., Schmitt, F., Edelman, R.R., Schomer, D.L., 1993. Monitoring the patient’s EEG during echo planar MRI. Electroencephalogr. Clin. Neurophysiol. 87. https://doi.org/10.1016/0013-4694(93)90156-P

Jorge, J., Bouloc, C., Bréchet, L., Michel, C.M., Gruetter, R., 2019. Investigating the variability of cardiac pulse artifacts across heartbeats in simultaneous EEG-fMRI recordings: A 7T study. Neuroimage 191, 21–35. https://doi.org/10.1016/j.neuroimage.2019.02.021

Jorge, J., Van der Zwaag, W., Figueiredo, P., 2014. EEG-fMRI integration for the study of human brain function. Neuroimage 102, 24–34. https://doi.org/10.1016/j.neuroimage.2013.05.114

Kirchner, W.K., 1958. Age differences in short-term retention of rapidly changing information. J. Exp. Psychol. 55, 352–358. https://doi.org/10.1037/h0043688

Kybic, J., Clerc, M., Faugeras, O., Keriven, R., Papadopoulo, T., 2006. Generalized head models for MEG/EEG: boundary element method beyond nested volumes. Phys. Med. Biol. 51, 1333–1346. https://doi.org/10.1088/0031-9155/51/5/021

Laufs, H., 2012. A personalized history of EEG-fMRI integration. Neuroimage. https://doi.org/10.1016/j.neuroimage.2012.01.039

Laufs, H., Daunizeau, J., Carmichael, D.W., Kleinschmidt, A., 2008. Recent advances in recording electrophysiological data simultaneously with magnetic resonance imaging. Neuroimage 40, 515–528. https://doi.org/10.1016/j.neuroimage.2007.11.039

Laufs, H., Kleinschmidt, A., Beyerle, A., Eger, E., Salek-Haddadi, A., Preibisch, C., Krakow, K., 2003. EEG-correlated fMRI of human alpha activity. Neuroimage 19, 1463–1476.

Lemieux, L., Salek-Haddadi, A., Josephs, O., Allen, P., Toms, N., Scott, C., Krakow, K., Turner, R., Fish, D.R., 2001. Event-related fMRI with simultaneous and continuous EEG: Description of the method and initial case report. Neuroimage 14. https://doi.org/10.1006/nimg.2001.0853

LeVan, P., Gotman, J., 2009. Independent component analysis as a model-free approach for the detection of BOLD changes related to epileptic spikes: A simulation study. Hum. Brain Mapp. 30. https://doi.org/10.1002/hbm.20647

LeVan, P., Maclaren, J., Herbst, M., Sostheim, R., Zaitsev, M., Hennig, J., 2013. Ballistocardiographic artifact removal from simultaneous EEG-fMRI using an optical motion-tracking system. Neuroimage 75, 1–11. https://doi.org/10.1016/j.neuroimage.2013.02.039

Lichstein, K.L., Riedel, B.W., Richman, S.L., 2000. The Mackworth Clock Test: A Computerized Version. J. Psychol. 134, 153–161. https://doi.org/10.1080/00223980009600858

Litvak, V., Eusebio, A., Jha, A., Oostenveld, R., Barnes, G.R., Penny, W.D., Zrinzo, L., Hariz, M.I., Limousin, P., Friston, K.J., Brown, P., 2010. Optimized beamforming for simultaneous MEG and intracranial local field potential recordings in deep brain stimulation patients. Neuroimage 50, 1578–1588. https://doi.org/10.1016/j.neuroimage.2009.12.115

Logothetis, N.K., Pauls, J., Augath, M., Trinath, T., Oeltermann, A., 2001. Neurophysiological investigation of the basis of the fMRI signal. Nature 412, 150–157. https://doi.org/10.1038/35084005

Loh, S., Lamond, N., Dorrian, J., Roach, G., Dawson, D., 2004. The validity of psychomotor vigilance tasks of less than 10-minute duration. Behav. Res. Methods, Instruments, Comput. 36, 339–346. https://doi.org/10.3758/BF03195580

Mandelkow, H., Halder, P., Boesiger, P., Brandeis, D., 2006. Synchronization facilitates removal of MRI artefacts from concurrent EEG recordings and increases usable bandwidth. Neuroimage 32, 1120–1126. https://doi.org/10.1016/j.neuroimage.2006.04.231

Mantini, D., Perrucci, M.G., Cugini, S., Ferretti, A., Romani, G.L., Del Gratta, C., 2007. Complete artifact removal for EEG recorded during continuous fMRI using independent component analysis. Neuroimage 34, 598–607. https://doi.org/10.1016/j.neuroimage.2006.09.037

Marino, M., Liu, Q., Koudelka, V., Porcaro, C., Hlinka, J., Wenderoth, N., Mantini, D., 2018. Adaptive optimal basis set for BCG artifact removal in simultaneous EEG-fMRI. Sci. Rep. 8, 8902. https://doi.org/10.1038/s41598-018-27187-6

Maris, E., 2012. Statistical testing in electrophysiological studies. Psychophysiology 49, 549–565. https://doi.org/10.1111/j.1469-8986.2011.01320.x

Maris, E., Oostenveld, R., 2007. Nonparametric statistical testing of EEG- and MEG-data. J. Neurosci. Methods 164, 177–190. https://doi.org/10.1016/j.jneumeth.2007.03.024

Marshall, T.R., Bergmann, T.O., Jensen, O., 2015. Frontoparietal Structural Connectivity Mediates the Top-Down Control of Neuronal Synchronization Associated with Selective Attention. PLOS Biol. 13, e1002272. https://doi.org/10.1371/journal.pbio.1002272

Masterton, R.A.J., Abbott, D.F., Fleming, S.W., Jackson, G.D., 2007. Measurement and reduction of motion and ballistocardiogram artefacts from simultaneous EEG and fMRI recordings. Neuroimage 37, 202–211. https://doi.org/10.1016/j.neuroimage.2007.02.060

Mayhew, S.D., Li, S., Kourtzi, Z., 2012. Learning acts on distinct processes for visual form perception in the human brain. J Neurosci 32, 775–786. https://doi.org/32/3/775 [pii] 10.1523/JNEUROSCI.2033-11.2012

Mayhew, S.D., Ostwald, D., Porcaro, C., Bagshaw, A.P., 2013. Spontaneous EEG alpha oscillation interacts with positive and negative BOLD responses in the visual-auditory cortices and default-mode network. Neuroimage 76, 362–372. https://doi.org/10.1016/j.neuroimage.2013.02.070

Mobascher, A., Brinkmeyer, J., Warbrick, T., Musso, F., Wittsack, H.J., Saleh, A., Schnitzler, A., Winterer, G., 2009. Laser-evoked potential P2 single-trial amplitudes covary with the fMRI BOLD response in the medial pain system and interconnected subcortical structures. Neuroimage 45, 917–926. https://doi.org/10.1016/j.neuroimage.2008.12.051

Moeller, F., Siniatchkin, M., Gotman, J., 2020. Simultaneous EEG and fMRI Recordings (EEG–fMRI), in: FMRI. Springer International Publishing, Cham, pp. 175–191. https://doi.org/10.1007/978-3-030-41874-8_13

Mullinger, K.J., Bowtell, R., 2011. Combining EEG and fMRI, in: Methods in Molecular Biology (Clifton, N.J.). pp. 303–326. https://doi.org/10.1007/978-1-61737-992-5_15

Mullinger, K.J., Brookes, M., Stevenson, C., Morgan, P., Bowtell, R., 2008a. Exploring the feasibility of simultaneous electroencephalography/functional magnetic resonance imaging at 7 T. Magn Reson Imaging 26, 968–977. https://doi.org/10.1016/j.mri.2008.02.014

Mullinger, K.J., Cherukara, M.T., Buxton, R.B., Francis, S.T., Mayhew, S.D., 2017. Post-stimulus fMRI and EEG responses: Evidence for a neuronal origin hypothesised to be inhibitory. Neuroimage 157, 388–399. https://doi.org/10.1016/j.neuroimage.2017.06.020

Mullinger, K.J., Mayhew, S.D., Bagshaw, A.P., Bowtell, R., Francis, S.T., 2014. Evidence that the negative BOLD response is neuronal in origin: a simultaneous EEG-BOLD-CBF study in humans. Neuroimage 94, 263–274.

Mullinger, K.J., Mayhew, S.D., Bagshaw, A.P., Bowtell, R., Francis, S.T., 2013. Poststimulus undershoots in cerebral blood flow and BOLD fMRI responses are modulated by poststimulus neuronal activity. Proc Natl Acad Sci U S A 110, 13636–13641. https://doi.org/10.1073/pnas.1221287110

Mullinger, K.J., Morgan, P.S., Bowtell, R.W., 2008b. Improved artifact correction for combined electroencephalography/functional MRI by means of synchronization and use of vectorcardiogram recordings. J Magn Reson Imaging 27, 607–616. https://doi.org/10.1002/jmri.21277

Mullinger, K.J., Yan, W.X., Bowtell, R., 2011. Reducing the gradient artefact in simultaneous EEG-fMRI by adjusting the subject’s axial position. Neuroimage 54, 1942–1950. https://doi.org/S1053-8119(10)01281-4 [pii] 10.1016/j.neuroimage.2010.09.079

Murta, T., Leal, A., Garrido, M.I., Figueiredo, P., 2012. Dynamic Causal Modelling of epileptic seizure propagation pathways: A combined EEG-fMRI study. Neuroimage 62. https://doi.org/10.1016/j.neuroimage.2012.05.053

Murta, T., Leite, M., Carmichael, D.W., Figueiredo, P., Lemieux, L., 2015. Electrophysiological correlates of the BOLD signal for EEG-informed fMRI. Hum. Brain Mapp. 36. https://doi.org/10.1002/hbm.22623

Muthukumaraswamy, S.D., 2010. Functional Properties of Human Primary Motor Cortex Gamma Oscillations. J. Neurophysiol. 104, 2873–2885. https://doi.org/10.1152/jn.00607.2010

Neugebauer, F., Möddel, G., Rampp, S., Burger, M., Wolters, C.H., 2017. The Effect of Head Model Simplification on Beamformer Source Localization. Front. Neurosci. 11. https://doi.org/10.3389/fnins.2017.00625

Niazy, R.K., Beckmann, C.F., Iannetti, G.D., Brady, J.M., Smith, S.M., 2005. Removal of FMRI environment artifacts from EEG data using optimal basis sets. Neuroimage 28, 720–737. https://doi.org/10.1016/j.neuroimage.2005.06.067

Nichols, T.E., Holmes, A.P., 2002. Nonparametric permutation tests for functional neuroimaging: A primer with examples. Hum. Brain Mapp. 15, 1–25. https://doi.org/10.1002/hbm.1058

Novitskiy, N., Ramautar, J.R., Vanderperren, K., De Vos, M., Mennes, M., Mijovic, B., Vanrumste, B., Stiers, P., Van den Bergh, B., Lagae, L., Sunaert, S., Van Huffel, S., Wagemans, J., 2011. The BOLD correlates of the visual P1 and N1 in single-trial analysis of simultaneous EEG-fMRI recordings during a spatial detection task. Neuroimage 54, 824–835. https://doi.org/10.1016/j.neuroimage.2010.09.041

Olbrich, S., Mulert, C., Karch, S., Trenner, M., Leicht, G., Pogarell, O., Hegerl, U., 2009. EEG-vigilance and BOLD effect during simultaneous EEG/fMRI measurement. Neuroimage 45, 319–332. https://doi.org/10.1016/j.neuroimage.2008.11.014

Oostendorp, T., van Oosterom, A., 1991. The potential distribution generated by surface electrodes in inhomogeneous volume conductors of arbitrary shape. IEEE Trans. Biomed. Eng. 38, 409–417. https://doi.org/10.1109/10.81559

Oostenveld, R., Fries, P., Maris, E., Schoffelen, J.M., 2011. FieldTrip: Open source software for advanced analysis of MEG, EEG, and invasive electrophysiological data. Comput Intell Neurosci 2011, 156869. https://doi.org/10.1155/2011/156869

Pakenham, D.O., Quinn, A.J., Fry, A., Francis, S.T., Woolrich, M.W., Brookes, M.J., Mullinger, K.J., 2020. Post-stimulus beta responses are modulated by task duration. Neuroimage 206, 116288. https://doi.org/10.1016/j.neuroimage.2019.116288

Pfurtscheller, G., Lopes da Silva, F.H., 1999. Event-related EEG/MEG synchronization and desynchronization: basic principles. Clin. Neurophysiol. 110, 1842–1857. https://doi.org/10.1016/S1388-2457(99)00141-8

Pfurtscheller, G., Stancák, A., Neuper, C., 1996. Post-movement beta synchronization. A correlate of an idling motor area? Electroencephalogr. Clin. Neurophysiol. 98, 281–293. https://doi.org/10.1016/0013-4694(95)00258-8

Quraan, M.A., Cheyne, D., 2010. Reconstruction of correlated brain activity with adaptive spatial filters in MEG. Neuroimage 49, 2387–2400. https://doi.org/10.1016/j.neuroimage.2009.10.012

Robinson, S.E., Vrba, J., 1999. Functional neuroimaging by synthetic aperture magnetometry (SAM), in: Yoshimoto, T., Kotani, M., Kuriki, S., Karibe, H., Nakasato, N. (Eds.), Recent Advances in Biomagnetism. Tohoku Univ. Press, Sendai, Japan, pp. 302–305.

Rothlübbers, S., Relvas, V., Leal, A., Murta, T., Lemieux, L., Figueiredo, P., 2015. Characterisation and Reduction of the EEG Artefact Caused by the Helium Cooling Pump in the MR Environment: Validation in Epilepsy Patient Data. Brain Topogr. 28, 208–220. https://doi.org/10.1007/s10548-014-0408-0

Russell, G.S., Jeffrey Eriksen, K., Poolman, P., Luu, P., Tucker, D.M., 2005. Geodesic photogrammetry for localizing sensor positions in dense-array EEG. Clin. Neurophysiol. 116, 1130–1140. https://doi.org/10.1016/j.clinph.2004.12.022

Scheeringa, R., Fries, P., Petersson, K.-M., Oostenveld, R., Grothe, I., Norris, D.G., Hagoort, P., Bastiaansen, M.C.M., 2011. Neuronal Dynamics Underlying High- and Low-Frequency EEG Oscillations Contribute Independently to the Human BOLD Signal. Neuron 69, 572–583. https://doi.org/10.1016/j.neuron.2010.11.044

Scheibe, C., Ullsperger, M., Sommer, W., Heekeren, H.R., 2010. Effects of parametrical and trial-to-trial variation in prior probability processing revealed by simultaneous electroencephalogram/functional magnetic resonance imaging. J Neurosci 30, 16709–16717. https://doi.org/30/49/16709 [pii] 10.1523/JNEUROSCI.3949-09.2010

Sekihara, K., Nagarajan, S.S., 2008. Adaptive Spatial Filters for Electromagnetic Brain Imaging, Series in Biomedical Engineering. Springer Berlin Heidelberg, Berlin, Heidelberg. https://doi.org/10.1007/978-3-540-79370-0

Sekihara, K., Nagarajan, S.S., Poeppel, D., Marantz, A., 2004. Asymptotic SNR of Scalar and Vector Minimum-Variance Beamformers for Neuromagnetic Source Reconstruction. IEEE Trans. Biomed. Eng. 51, 1726–1734. https://doi.org/10.1109/TBME.2004.827926

Sekihara, K., Nagarajan, S.S., Poeppel, D., Marantz, A., Miyashita, Y., 2001. Reconstructing spatio-temporal activities of neural sources using an MEG vector beamformer technique. IEEE Trans. Biomed. Eng. 48, 760–771. https://doi.org/10.1109/10.930901

Srivastava, G., Crottaz-Herbette, S., Lau, K.M., Glover, G.H., Menon, V., 2005. ICA-based procedures for removing ballistocardiogram artifacts from EEG data acquired in the MRI scanner. Neuroimage 24, 50–60. https://doi.org/10.1016/j.neuroimage.2004.09.041

Sweet, L.H., 2011. N-Back Paradigm, in: Encyclopedia of Clinical Neuropsychology. Springer New York, New York, NY, pp. 1718–1719. https://doi.org/10.1007/978-0-387-79948-3_1315

Takemi, M., Masakado, Y., Liu, M., Ushiba, J., 2013. Event-related desynchronization reflects downregulation of intracortical inhibition in human primary motor cortex. J. Neurophysiol. 110, 1158–1166. https://doi.org/10.1152/jn.01092.2012

Tyvaert, L., LeVan, P., Grova, C., Dubeau, F., Gotman, J., 2008. Effects of fluctuating physiological rhythms during prolonged EEG-fMRI studies. Clin. Neurophysiol. 119. https://doi.org/10.1016/j.clinph.2008.07.284

Uji, M., Jentzsch, I., Redburn, J., Vishwanath, D., 2019. Dissociating neural activity associated with the subjective phenomenology of monocular stereopsis: An EEG study. Neuropsychologia 129, 357–371. https://doi.org/10.1016/j.neuropsychologia.2019.04.017

Uji, M., Wilson, R., Francis, S.T., Mullinger, K.J., Mayhew, S.D., 2018. Exploring the advantages of multiband fMRI with simultaneous EEG to investigate coupling between gamma frequency neural activity and the BOLD response in humans. Hum. Brain Mapp. 39, 1673–1687. https://doi.org/10.1002/hbm.23943

van Drongelen, W., Yuchtman, M., Van Veen, B.D., van Huffelen, A.C., 1996. A spatial filtering technique to detect and localize multiple sources in the brain. Brain Topogr. 9, 39–49. https://doi.org/10.1007/BF01191641

van Veen, B.D., Buckley, K.M., 1988. Beamforming: a versatile approach to spatial filtering. IEEE ASSP Mag. 5, 4–24. https://doi.org/10.1109/53.665

van Veen, B.D., van Drongelen, W., Yuchtman, M., Suzuki, A., 1997. Localization of brain electrical activity via linearly constrained minimum variance spatial filtering. Biomedical 44, 867–880. https://doi.org/10.1109/10.623056

Vanderperren, K., De Vos, M., Ramautar, J.R., Novitskiy, N., Mennes, M., Assecondi, S., Vanrumste, B., Stiers, P., Van den Bergh, B.R.H., Wagemans, J., Lagae, L., Sunaert, S., Van Huffel, S., 2010. Removal of BCG artifacts from EEG recordings inside the MR scanner: A comparison of methodological and validation-related aspects. Neuroimage 50, 920–934. https://doi.org/10.1016/j.neuroimage.2010.01.010

Vanderperren, K., Ramautar, J., Novitskiy, N., De Vos, M., Mennes, M., Vanrumste, B., Stiers, P., Van den Bergh, B.R.H., Wagemans, J., Lagae, L., Sunaert, S., Van Huffel, S., 2007. Ballistocardiogram artifacts in simultaneous EEG-fMRI acquisitions. Int. J. Bioelectromagn. 9, 146–150.

Wilson, R., Mullinger, K.J., Francis, S.T., Mayhew, S.D., 2019. The relationship between negative BOLD responses and ERS and ERD of alpha/beta oscillations in visual and motor cortex. Neuroimage 199, 635–650. https://doi.org/10.1016/j.neuroimage.2019.06.009

Xia, H., Ruan, D., Cohen, M.S., 2014. Removing ballistocardiogram (BCG) artifact from full-scalp EEG acquired inside the MR scanner with Orthogonal Matching Pursuit (OMP). Front. Neurosci. https://doi.org/10.3389/fnins.2014.00218

Xia, H., Ruan, D., Cohen, M.S., 2013. Coupled basis learning and regularized reconstruction for BCG artifact removal in simultaneous EEG-FMRI studies, in: Proceedings – International Symposium on Biomedical Imaging. https://doi.org/10.1109/ISBI.2013.6556642

